# Interpretable Machine Learning Model of Receptor Dynamics Reveals AT1R Allostery and a Negative Allosteric Modulator

**DOI:** 10.64898/2026.08.01.742107

**Authors:** Hanyu Chen, Yoon Namkung, Zahra Asadi Jafari, Wijnand J. C. van der Velden, Elizaveta Mukhaleva, Grady Kestler, Andrei S. Rodin, Sergio Branciamore, Stephane A. Laporte, Nagarajan Vaidehi

## Abstract

Allosteric modulation of G protein–coupled receptors (GPCRs) offers major advantages in receptor selectivity and signaling control; yet systematic approaches to identify allosteric modulators, define their binding sites, and map the underlying allosteric networks remain limited. Current molecular dynamics (MD) and machine learning (ML)-based methods often rely on correlation-driven or black-box models that provide limited mechanistic insight. We developed an interpretable probabilistic framework that extracts residue-level dependencies from MD ensembles using Bayesian network modeling (BNM). By representing each residue through its local interaction energy, BNM identifies both local and long-range energetic couplings and maps the allosteric communication pathways linking the AngII binding site to the G-protein interface in the angiotensin II type 1 receptor (AT1R). To functionally prioritize these pathways, we integrated BNM with comprehensive mutational analysis, combining whole-receptor alanine mutagenesis data with exhaustive in silico deep mutational scanning to validate BNM-predicted hotspots. This approach recovered state-dependent allosteric communities, revealed residues in noncanonical regions that regulate Gα_q_ coupling and identified positions whose functional importance emerged only with specific, predicted substitutions, as well as highlighted a cryptic intracellular pocket enriched in communication hubs. Guided by these network-derived residues and pocket geometries, structure-based virtual screening identified a small, fragment-like molecule negative allosteric modulator (NAM) named Q2 that attenuates AngII-mediated Gα_q_ signaling. Mutational mapping supports Q2 binding adjacent to the G-protein interface, consistent with its mechanism of action. Together, these results establish a generalizable and interpretable framework for uncovering GPCR allosteric communication networks and discovering modulators that exploit these networks.

## Introduction

G protein–coupled receptors (GPCRs) mediate diverse physiological processes and remain major therapeutic targets^1–3^. Their ability to interconvert among multiple conformational states enables bidirectional communication between the ligand binding pocket and the intracellular G protein coupling interface (GPI), forming an allosteric communication network that is spread across different structural regions of the receptor^4–10^. This dynamic communication creates opportunities for selective pharmacology, as small molecules binding outside the orthosteric site can modulate signaling with improved subtype specificity^11,12^. Yet the allosteric pockets that confer such advantages are often cryptic, transient, state-dependent, and not readily apparent from static structures ^13,14^, making their systematic identification challenging.

Molecular dynamics (MD) simulations capture the allostery and cryptic binding pocket formation embedded in the dynamics by generating ensembles of GPCR conformations in active and inactive states. However, analyzing high-dimensional MD trajectories in a data-driven fashion is a critical first step to developing a systematic method for identifying allosteric communication networks and further allosteric binding sites in GPCRs. Traditional analyses, such as principal component analysis (PCA) and correlation-based dynamic networks, highlight collective motions or contact fluctuations but do not distinguish direct from transitive relationships^15–17^. Consequently, these approaches often amplify local correlations at the expense of identifying residues that genuinely drive long-range allostery. Similarly, MD based mapping of cryptic cavities reveals where pockets appear, but not whether they participate in functionally relevant allosteric communication with the orthosteric or G protein interface (GPI) sites^14,18,19^.

Machine learning (ML) approaches have begun to address some of these limitations. Early conventional classifiers and more recent deep learning models trained on structures or sequences can identify residues associated with allosteric regulation^20–25^. However, these tools remain largely correlational with limited interpretability and often rely on static inputs or pretraining that may not be well matched to the system under study. Without capturing conditional dependencies across receptor conformational states, such models cannot determine how specific residues modulate information flow in different signaling states. Thus, although MD and ML each yield useful insights, neither approach alone provides a robust framework for inferring the mechanistic architecture of allostery.

To fill this gap, we apply Bayesian Network Modeling (BNM), a probabilistic Interpretable unsupervised ML method, to MD ensembles to infer conditional probabilistic dependencies among residues^26^. By representing each residue through its interaction energy with the surrounding environment, the method learns probabilistic influence pathways, filters out spurious correlations, and quantifies uncertainty^27–29^. This network centric, interpretable framework yields directed acyclic graphs that reveal how information propagates through the receptor structure. In this work, we further develop a Differential BNM (DBNM) analysis to rigorously compare networks recovered from distinct receptor states, enabling the identification of residues whose communication roles are reorganized in different functional states, such as inactive and active states of GPCRs (Fig. 1a). This contrasts with existing methods by directly mapping state-dependent changes onto explicit mechanistic influence patterns, rather than merely associating them with correlated motions.

**Figure 1.**
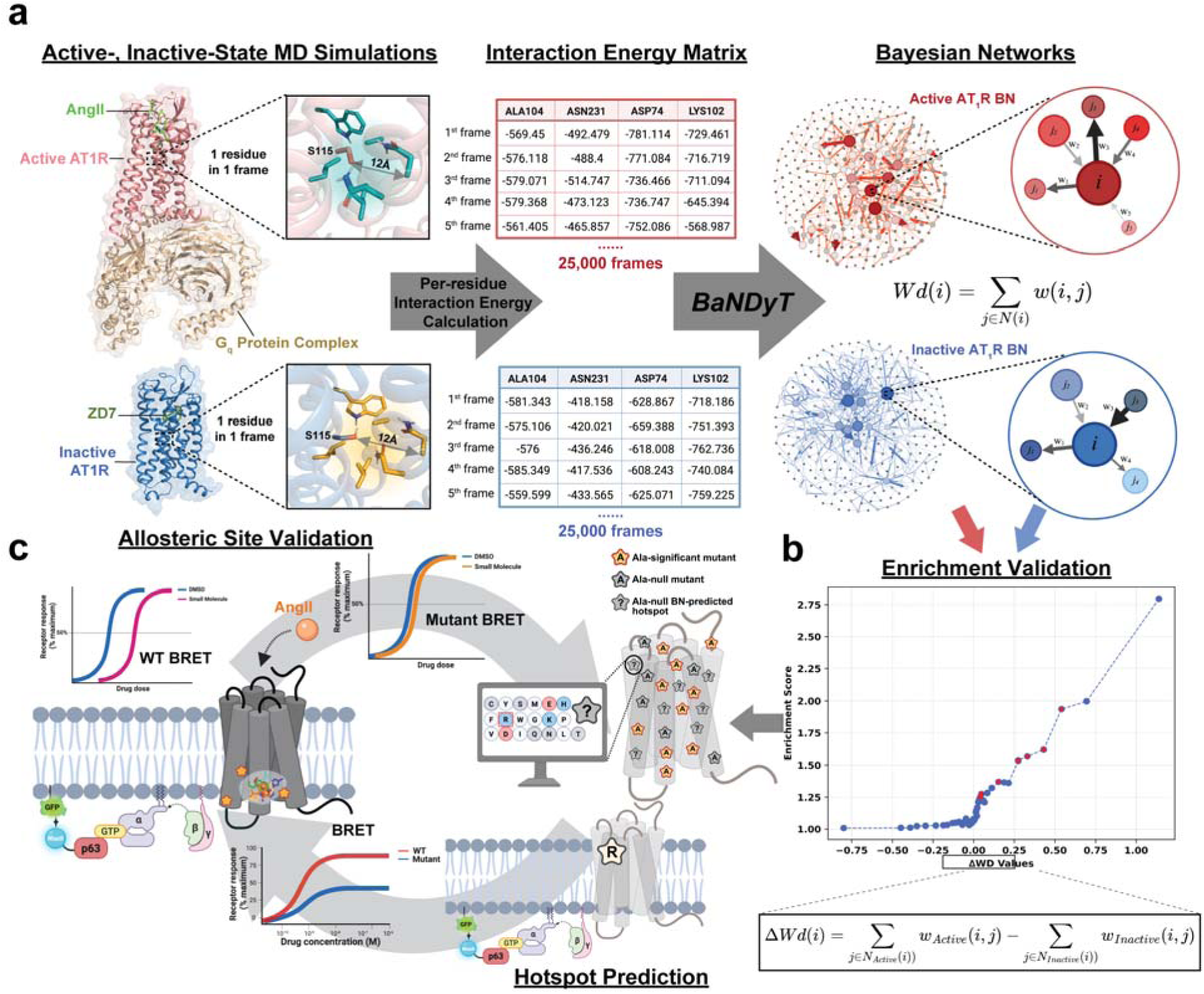
Workflow of Bayesian network analysis and the BRET experimental assays. (**a**) MD simulations were done on fully active (AngII:AT1R: Gq) and antagonist ZD7 bound inactive (AT1R) states. The residue interaction energies were calculated from five trajectories (25,000 MD snapshots) and used as input variables to generate state-specific Bayesian networks for the active (red) and inactive (blue) states. The difference in the weighted degree for each node (residue) was calculated (Δwd_i_), and residues were ranked accordingly. (**b**) Validation of the calculated DWd for ranked residues from Bayesian networks using enrichment analysis with the experimental positives from alanine scanning mutagenesis. Enrichment curves were generated by calculating, at each rank threshold, the score for experimental positives relative to background, with statistical significance assessed using Fisher’s exact test (red points indicate p < 0.05). (**c**) Iterative-predict-test cycle: Predicted non-alanine mutations for high ranking ΔWd residue positions that did not show effect upon alanine mutation, were tested for effects on AngII-mediated Gq activation. Gq BRET biosensor (Gα_q_, P63-RlucII, and rGFP-CAAX) assay used for generating dose–response curves to assess receptor activation in wild-type and mutant AT1R. Also shown is the procedure to predict and test small fragment-like molecules to modulate AngII mediated AT1R activation. Together, these assays show the iterative workflow used to predict residues that are hotspots in allosteric communication pathways from MD and Bayesian network modeling (BNM) analysis and to test them using BRET assays for experimental validation of allosteric sites.

We apply our framework to the angiotensin II type 1 receptor (AT1R), a prototypical class A GPCR, activated by angiotensin II (AngII) and central to cardiovascular regulation ^30–32^. Although high-resolution structures have captured snapshots of AT1R in active and inactive states ^33,34^, the dynamic communication that links the ligand-binding pocket, intracellular interface, and cryptic allosteric sites remains incompletely understood. By combining extensive MD simulations with DBNM-based analysis, whole-receptor alanine scanning data and mutants’ Gα_q_ functionality, we identified residue communities in AT1R that form contiguous allosteric communication pathways between functional regions. We identified cryptic small molecule binding pockets in the intracellular region of the receptor that was enriched in residues predicted to regulate Gα_q_ coupling. Virtual ligand screening of one of these pockets led to the discovery of a small, fragment-like molecule negative allosteric modulator (NAM), Q2, which binds allosteric to AngII and above the GPI and inhibits AngII-mediated signaling. Our findings demonstrate that a dynamics-driven, interpretable probabilistic ML framework allows mapping the causal architecture of GPCR allostery and guide the rational discovery of small molecule allosteric modulators

## Results

### Alanine-scanning mutagenesis effects on AT1R Gα_q_ coupling

Previously, we generated a library of 359 AT1R alanine mutants or glycine in the case of native alanine, to identify residues critical for G protein coupling, using the downstream PKC-c1b BRET biosensors (Fig. 1) that report endogenous Gα_q/11_ activation^35^. We determined the pharmacological EC_50_ and Emax values of AngII across mutants using dose–response curves, and receptor expression levels were titrated to define the minimum threshold required for maximal G_q/11_ response. Mutants expressing less than half the wild-type (WT) receptor level (30 in total) were excluded from analysis. The resulting dataset, which provided a comprehensive map of residues contributing to Gα_q_ coupling, was used to validate DBNM applied to MD simulation trajectories of AT1R.

### Bayesian network models of active and inactive AT1R ensembles

Starting from the respective crystal structures, we performed all-atom MD simulations on the fully active state of AT1R bound to AngII and GαqGβγ (pdb ID: 7F6G) and the antagonist (ZD7) bound inactive state of AT1R (pdb ID: 4YAY) in a POPC lipid bilayer (see Methods for details). For each conformational state, we performed five MD runs each 1 μs long followed by BNM analysis using our software BaNDyT^27^. We used the nonbonded interaction energy between each residue and its surrounding residues within 12Å radius in AT1R, calculated for every MD snapshot, as input variables to generate the BN model (Fig. 1a) for both the inactive and active states of AT1R. In the resulting BN, AT1R residues are the nodes, the edges connecting two nodes show correlated movement or co-dependencies in residue dynamics and edge weights are the probability of the co-dependencies.

The distribution of residue interaction energies used as inputs to the BNM is shown in Supplementary Fig. 1A, B, revealing that residues in the intracellular loops (ICLs) display greater energy spread than residues in the transmembrane (TM) helices in both the active and inactive states. As inputs to BaNDyT, we discretized the continuous residue interaction energy using a maximum-entropy binning procedure^36^, which yields an approximately equal-frequency partition with minimal distributional assumptions. Eight bins were used (see Methods for details). We fitted separate BN models for the active and inactive states of AT1R (Fig. 1). For each residue node, the weighted degree (Wd) which is the sum of the strengths of all incident edges, quantifies its aggregated co-dependencies with neighboring nodes in the probabilistic space, irrespective of their spatial proximity. To evaluate the statistical robustness of the Wd, we performed a bootstrap procedure in which the original MD-derived interaction energy matrix was resampled 1,000 times and mutual information was recomputed for each resampled dataset. These mutual information values were used to reweight the fixed BN topology for every replicate, yielding a distribution of Wd values per residue. Wd estimates were highly consistent across bootstrap samples in both signaling states, and the resulting differential weighted degree (ΔWd) showed reproducible state-dependent shifts (Supplementary Fig. 2). These results indicate that the allosteric communication patterns encoded in the BN are stable with respect to sampling variability in the underlying MD ensemble.

### Bayesian network modeling of AT1R dynamics reveals state-dependent information flow

To compare information flow within each AT1R conformational state, we calculated the Wd of each residue from the BNM of the active and inactive AT1R ensembles and performed a quartile analysis of Wd. Respectively in active- and inactive-state BNMs, residues in the top quartile of Wd were grouped into three functional regions: ligand-binding site (LBS), Gα_q_ protein interface (GPI), and Mediators (residues that are not in LBS or GPI) to quantify how communication is distributed across the receptor. Network edges (representing co-dependencies in the dynamics) connecting residues at a distance in the BNM (>10 Å separation between the Cα atoms) were classified as allosteric. In the inactive state, residues in the top quartile of Wd, exhibit co-dependencies primarily within the GPI region (33%) and between GPI and mediator residues (29%), with limited coordinated motion observed within the LBS (3%) and mediator domains (18%) (Fig. 2a). The higher co-dependencies within the GPI region indicate the more coordinated motion among the intracellular part of the receptor to keep the Gαq protein binding cavity from opening in the inactive state (Fig. 2a, b). In the active-state AngII–Gα_q_ signaling complex, the co-dependencies within the GPI decrease showing that the intracellular interface has either decreased coordinated motion or no motion at all; and the mediator residues emerge as a central hub. Notably, strong interdomain links between Mediators and LBS (28%) and between Mediators and GPI (16%) indicate extensive allosteric integration (Fig. 2d). Our comparison contrasts the allosteric co-dependencies or allosteric communication mechanisms of an active AngII–Gα_q_ complex with an antagonist bound receptor devoid of G protein.

**Figure 2.**
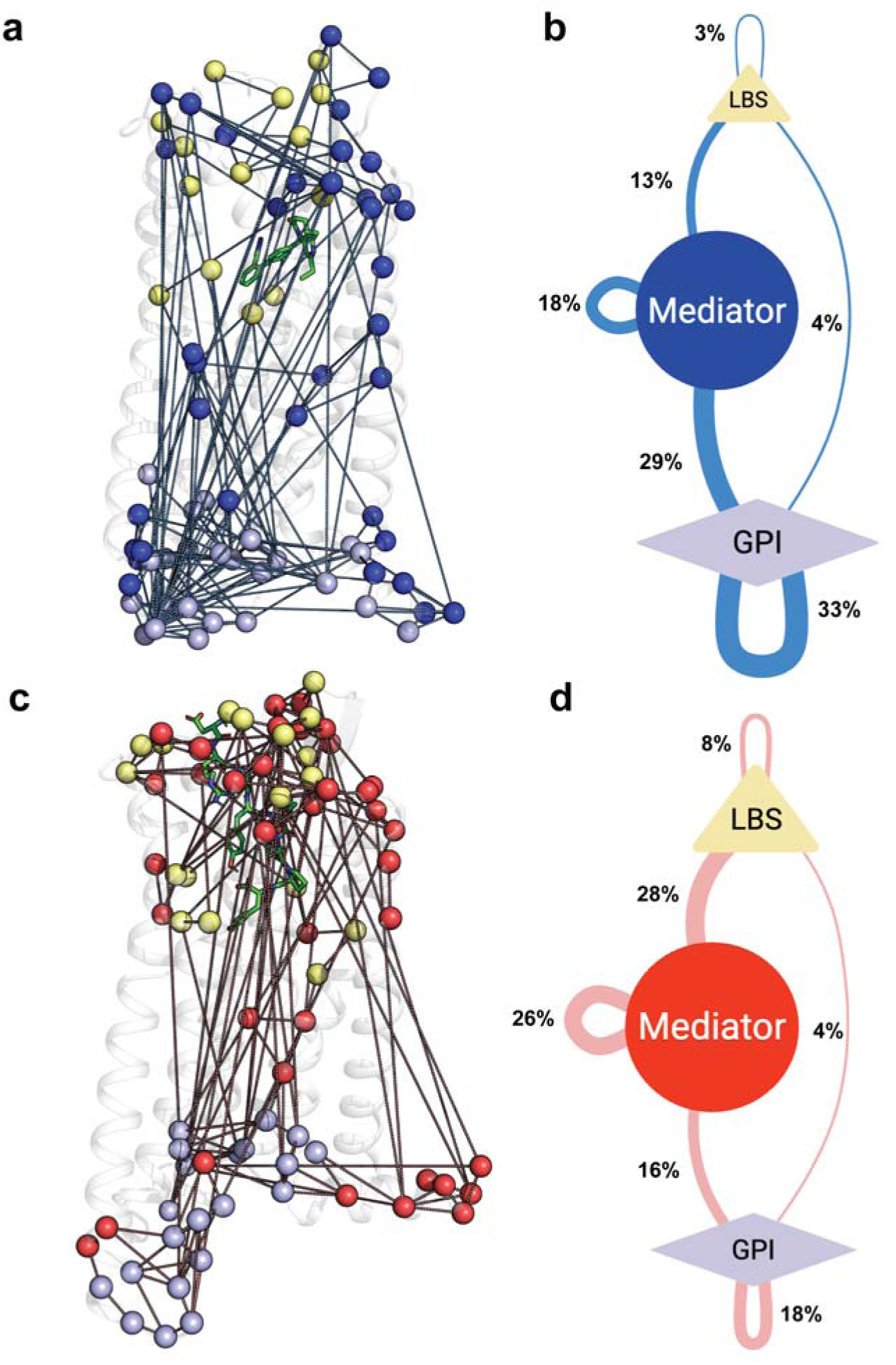
Mechanism of Information flow between extracellular and intracellular regions in inactive and active states of AT1R. (**a**) The Bayesian network edges among residues that rank in the top quartile by Wd in the inactive AT1R state shown on the structure of the inactive state of AT1R (PDB ID: 4YAY). Colored spheres represent residues by functional regions: ligand-binding site (LBS, yellow), mediator region (blue) and Gαq protein interfacing (GPI) residues (purple) as identified by residue contact analysis in the AT1R: Gα_q_ interface from MD simulations of the active state. (**b**) Summary of the pattern of allosteric communication between different structural regions of AT1R in the inactive state. The thickness of the blue lines connecting the different structural regions is proportionate to the sum of all the edge strengths between residue pairs located in each of the three structural regions of AT1R. The shape sizes are proportionate to the number of residues in each group in the top quartile of Wd. (**c**) The Bayesian network edges among residues that rank in the top quartile by Wd in the active AT1R state shown on the structure of the active state of AT1R (PDB ID: 7F6G). Colored spheres represent residues by functional regions: ligand-binding site (LBS, yellow), mediator region (red) and Gα_q_ protein interfacing (GPI) residues (purple) as identified by residue contact analysis in the AT1R: Gα_q_ interface from active state MD simulations. (**d**) Summary of the pattern of allosteric communication between different structural regions of AT1R in the active state. The thickness of the pink lines connecting the different structural regions is proportionate to the sum of all the edge strengths between residue pairs located in each of the three structural regions of AT1R. The shape sizes are proportionate to the number of residues in each group in the top quartile of Wd in the active state of AT1R.

Consistent with the redistribution of top-quartile Wd connectivity, visualization of allosteric (>10 Å) dependencies reveals a pronounced reorganization of global communication across receptor states. In the inactive ensemble, direct and allosteric co-dependencies are diffused and fragmented across the intracellular face, lacking a dominant communication axis (Supplementary Fig. 3A). In contrast, the active state exhibits a coherent network of allosteric dependencies spanning multiple transmembrane helices and converging on the intracellular signaling interface (Supplementary Fig. 3B), forming a continuous allosteric communication spine linking distal receptor domains. Together, these observations indicate a state-dependent transition from locally constrained, regionally isolated motions to a globally integrated allosteric architecture that supports efficient transmission of ligand-induced conformational changes to the G protein–binding interface in AngII–Gα_q_ complex. Although receptor activation and G protein engagement cannot be fully disentangled here, this reorganization encodes functionally relevant information flow and provides a structural framework for interpreting residue-level network metrics.

### Differential weighted degree from BNM recapitulates the residues that regulate G**α**_q_ coupling to AT1R

On this basis, we asked whether state-dependent changes in Wd could predict functional sensitivity of each residue on AT1R:Gα_q_ coupling. For each residue we computed the difference in Wd extracted from the BNM of the active and inactive states (ΔWd) (Fig. 1a). We then compared ΔWd rankings with experimental outcomes from a whole-receptor alanine scan, where functional impact was quantified as the change in relative activity (RA) for AT1R:Gα_q_ signaling (Δlog(Emax/EC_50_)) compared to WT^35^. We found that residues with higher positive ΔWd, indicating increased probabilistic co-dependencies in the active state, were significantly enriched among functionally impactful alanine mutants (Fig. 1b). The enrichment scores were calculated as the fraction of experimental positives within the top-*n* ranked residues, normalized to the overall fraction of positives in the dataset (random expectation, ES = 1). This enrichment driven by the differential (state-contrast) Wd outperformed rankings based on Wd in either state alone (Supplementary Fig. 4C-F), demonstrating that ΔWd captures state-dependent predictive information relevant to Gα_q_ coupling. Residues ranked by either their positive ΔWd (greater importance in the receptor’s active state) or negative ΔWd (greater importance in the receptor’s inactive state) both correlated with experimental outcomes (Supplementary Fig. 4A, B). Thus, ΔWd provides a quantitative criterion for ranking residues in the receptor by their likelihood of functionally regulating Gα_q_ coupling.

Residues that showed maximum differences in correlated movements between receptor’s active and inactive states are shown in a waterfall plot (Fig. 3a). Residue G22^1.26^, for instance, showed significantly higher co-dependency with both spatially neighboring and allosteric residues in the active state compared to the inactive state (Fig. 3b). R126^3.50^ showed low ΔWd values which stems from its high weighted degree in both conformational states (Fig. 3c). In contrast, residue F250^6.45^ next to the PIF motif (Ile^3.40^, Pro^5.50^ and Phe^6.44^) and part of the TM5/TM6 aromatic residue cluster (F208^5.51^/F249^6.44^/F250^6.45^) that was found to distinguish between inactive and active AT1R conformation states ^37,38^, exhibited a negative ΔWd, reflecting reduced network connectivity in the active state relative to the inactive state (Fig. 3d).

**Figure 3.**
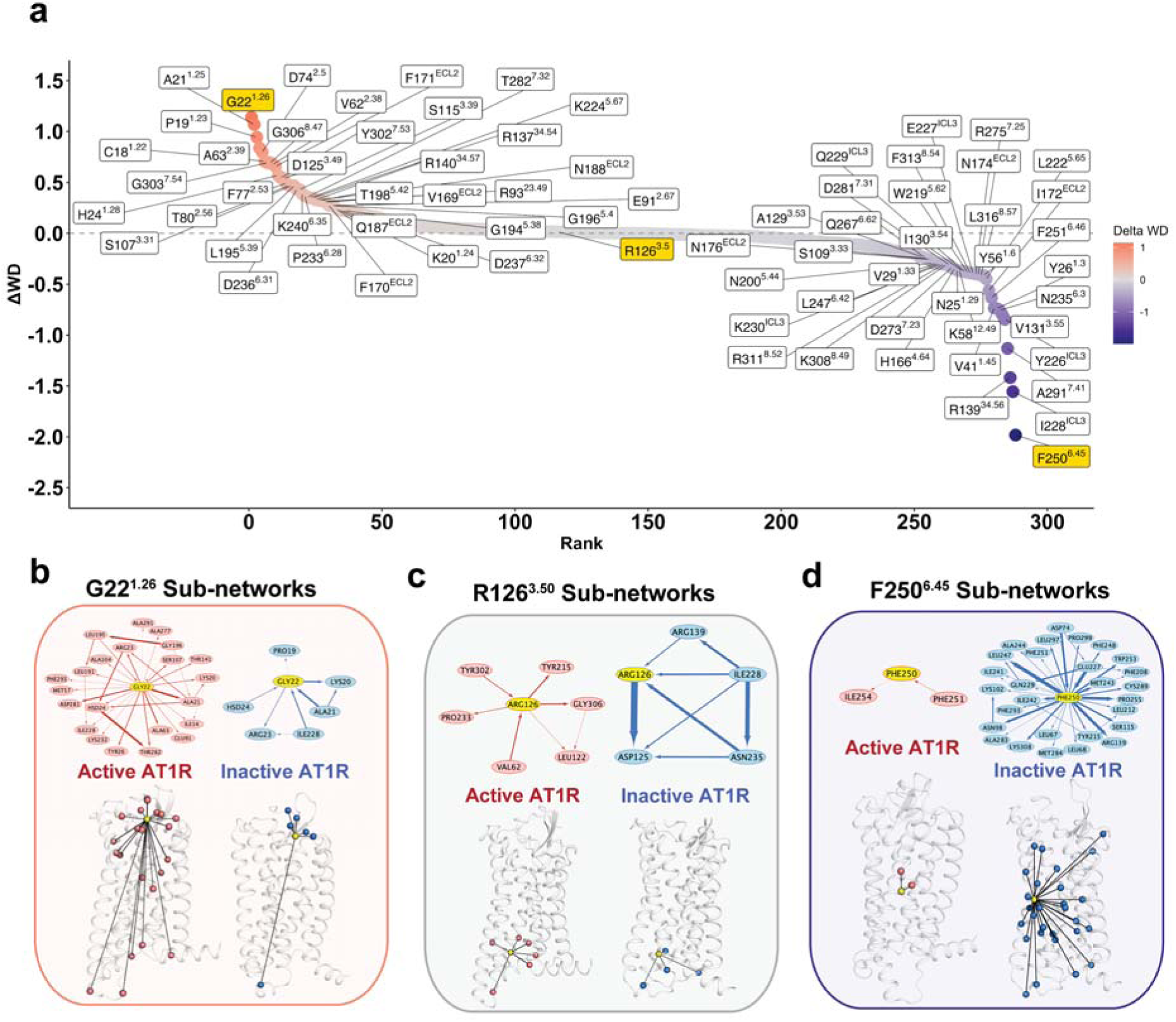
Weighted degree difference between active and inactive states follows the power law. (**a**) Waterfall plot showing the ranked ΔWd of all 289 residues common to active (PDB ID: 7F6G) and inactive (PDB ID: 4YAY) state AT1R crystal structures. (**b**) Sub-network of Gly22^1.26^ with a high positive ΔWd shown in active and inactive states. (**c**) Sub-network of Phe250^6.45^ with high negative ΔWd shown in active and inactive states. (**d**) Sub-network of Arg126^3.50^ with ΔWd close to zero in active and inactive states.

### Hierarchical clustering reveals residue communities involved in allosteric communication and targetable for allosteric modulator identification

Having shown that ΔWd captures state-dependent rewiring of residue-level subnetworks (Fig. 3), we next asked whether residues with extreme ΔWd values assemble into spatially coherent communities across the receptor. Because high ΔWd values highlight residues whose network influence changes between active and inactive states, we performed quartile-based analyses on residues with positive and negative ΔWd values and applied hierarchical clustering based on pairwise Cα–Cα distances in the corresponding structures. Residues in each ΔWd quartile were mapped onto the receptor structures, and the resulting distance matrices were hierarchically clustered to assess whether high-ranking residues are spatially adjacent or distributed across the receptor (Supplementary Fig. 5).

In the active state, residues in the top quartile of positive ΔWd, reflecting increased connectivity relative to the inactive state, form continuous clusters spanning the LBS, mediator region, and GPI, indicating strong allosteric coupling across these functional domains (Supplementary Fig. 5A). In contrast, residues in the top quartile of negative ΔWd form more compartmentalized clusters, enriched near the inner core and intracellular interface of AT1R with fewer continuous links between the LBS, mediator, and GPI (Supplementary Fig. 5I), indicating a rewired communication pattern that favors stabilization of the inactive ensemble. These contrasting connectivity patterns emphasize the mediator region’s central role in dynamically coupling functional sites in AT1R and highlight its potential as a strategic target for therapeutic intervention. Lower-ranked ΔWd quartiles show progressively diffuse spatial distributions, lacking the coherent communities observed among top-ranking residues (Supplementary Fig. 5B-D and 5F-H for positive ΔWd; Supplementary Fig. 5J-L and 5N-P for negative ΔWd).

### Predicted top-scoring residues from DBNM show functional effects upon non-alanine mutations

Together, quartile and clustering analyses revealed that residues with high positive ΔWd values form coherent allosteric communities enriched for functional importance (Fig. 1). However, among the 36 residues in the top ΔWd quartile, 16 showed no significant effects on AT1R–Gα_q_ coupling upon alanine/glycine substitution in the receptor ^35^. These positions may represent false negatives of alanine scanning, or alternatively, may require specific side-chain mutations to reveal their more penetrant functional role in AT1R. To distinguish between these possibilities, we performed an *in silico* deep mutational scan at each of the 16 residue positions, systematically evaluating all 20 amino acid substitutions in each position. For each mutant, we computed changes between mutant and the WT in interaction energies between the AT1R and Gα_q_ protein (ΔΔG_affinity) and in total energy of the AT1R: Gα_q_ system (ΔΔG_stability) (Data S2). Non-alanine mutations were selected for experimental testing based on their computed change on Gα_q_ interaction energy with AT1R and stability, and preservation of key structural interactions such as number of hydrogen bonds and van der Waals interaction (see Methods). We predicted more than one non-alanine mutations in some of the residue positions. Experimental testing of 26 predicted non-alanine mutations across 16 residue positions revealed significant functional effects (EC_50_ or Emax and/or RA) on AT1R-Gα_q_ signaling at 10 residue positions (Fig. 4 and Supplementary Table 1). Specifically, non-alanine residue mutations significantly altered Gα_q_ activation as revealed by changes in Emax (N188I^ECL2^, T198H^5.42^, P233L^6.28^, D236R^6.31^, and K240P^6.35^), EC_50_ (V169W^ECL2^ and H24P^1.28^) or both parameters (A21W^1.25^, S107P^3.31^, and L195E^5.39^), when compared to the WT.

**Figure 4.**
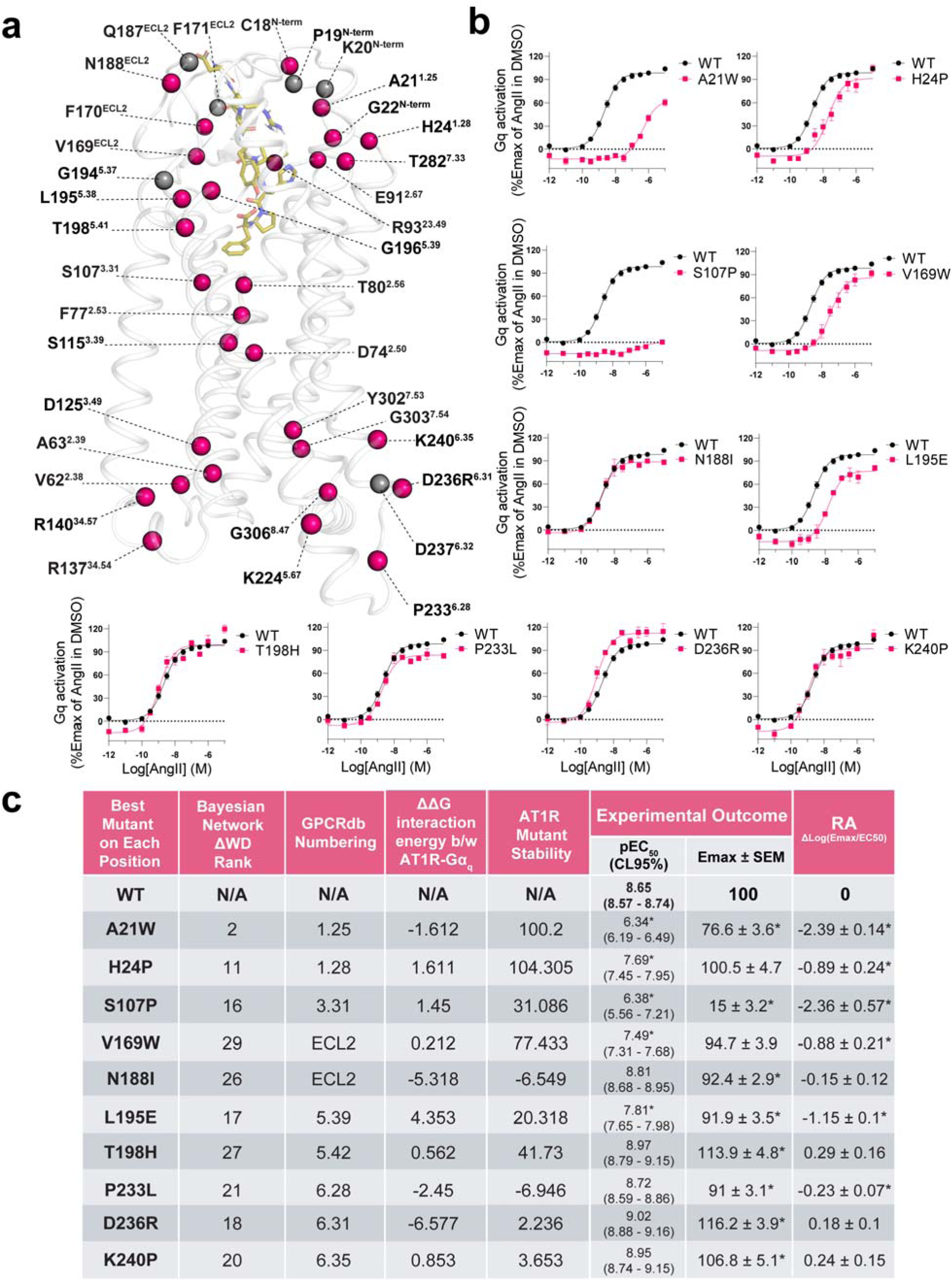
Experimental validation of BN-predicted high scoring residues with non-alanine mutations. (**a**) Structural visualization of residues ranked in the top quartile by ΔWd by BNM analysis (Data S1). Colored spheres indicate prediction and testing outcomes: pink = true positives (predicted and tested by alanine or non-alanine mutation(s), gray = tested with non-alanine mutation and have not shown a significant effect. (**b**) Dose–response curves using the Gq BRET biosensor (Gα_q_, P63-RlucII, and rGFP-caax) for predicted non-alanine mutations at high-ranking ΔWD residue positions that showed no significant effect with alanine substitution. Data are normalized to WT, and error bars indicate SEM from three biological replicates (also see Supplementary Fig.6 in Supporting Information for data on all mutants). (**c**) Table of the non-alanine mutations predicted by a computational deep mutational scan. 3^rd^ column contains the residue-based numbering system used for GPCRs. 4^th^ column is the difference in the interaction energy of the Gα_q_ protein with AT1R in the mutant versus the wild type (WT). The difference in total energy of the mutants versus the WT, the mutants’ stability is shown in 5^th^ column. More details in the Methods section. The 6^th^ and 7^th^ columns of the table are the changes in EC_50_ and Emax for each mutation in response to AngII-mediated AT1R activation, as measured by BRET.

Although most predicted non-alanine substitutions at high-ΔWd sites showed changes in AT1R-Gα_q_ coupling, six positions (P19H^1.23^, K20D^1.24^, F171D^ECL2^, Q187W ^ECL2^, G194H ^5.38^, D237G^6.32^) remained functionally indistinguishable from WT. This absence of a measurable effect is nonetheless compatible with the BNM-derived rankings and aligns with several mechanistic considerations. First, four of these residue positions fall near the lower boundary of the first positive ΔWd quartile (Fig. 3a, Supplementary Data 1), close to the selection cutoff, and would therefore be expected to exert relatively modest influence compared with higher-ranked positions. Additionally, although these residues were scored with high ΔWd values and occupy central positions in the BNM-derived information-flow network, it is possible that there are redundant allosteric communication pathways across the ensemble and hence the pathways going through these residues do not individually control the output. This is evident from the dense communication network shown in Supplementary Fig. 3. The silent predicted non alanine mutants likely reflect network redundancy and mutation choice rather than false-positive predictions. We have observed such redundancy in our recent work on AT1R and prostaglandin FP receptors coupling to different G proteins described in an accompanying manuscript (Boora *et al.*, submitted). Taken together, our DBNM analysis expands the mutational coverage of key allosteric residue communities by revealing positions whose functional roles are masked by conservative substitutions.

In a previous study we showed that in addition to residues within the ligand-binding pocket, alanine mutations at distal positions from the orthosteric site, like M134^34.51^, T141^4.38^, F301^7.52^, and F309^8.50^ which are in the intracellular region of AT1R (Fig. 5), decreased AT1R-mediated Gα_q_ response as well as binding affinity of AngII to AT1R ^35^. Radiolabeled AngII binding experiments showed that T141A ^4.38^ resulted in 75% loss in AngII binding compared to the WT. Mutants M134A ^34.51^, F301A ^7.52^, and F309A ^8.50^ showed AngII binding loss by 50%, 100%, and 100% respectively (Supplementary Fig. 10 in ^35^). In our current BNM analysis each of these residues showed strong allosteric co-dependencies or correlated movement with residues directly in the AngII binding site or in the second shell of AngII binding site (Fig. 5). Specifically, T141^4.38^ showed strong correlated motion allosterically (high edge weight in BNM connecting T141^4.38^ and G22^N-term^) with G22^N-term^, which impaired AngII binding when mutated to Ala ^35^; M134^34.51^ showed correlated movement with ligand binding site residue N188^ECL2^ (as identified from MD simulations of the active state of AT1R); F301^7.52^ and F309^8.50^ both strongly co-dependent with nearby G306^8.47^, which allosterically showed correlated motion with A21^N-term^. Despite not contacting AngII, A21^N-term^ is in proximity of the ligand, and has a high positive ΔWd with mutation to Trp impaired both LogEC_50_ and Emax (Fig. 4b). Through these allosteric co-dependencies observed in the BNM, mutations at M134^34.51^, T141^4.38^, F301^7.52^, and F309^8.50^ could destabilize AngII–AT1R interactions indirectly by disrupting allosteric co-dependency. Thus, BNM unifies prior mutational observations within a coherent, network-based framework of AT1R allostery.

**Figure 5.**
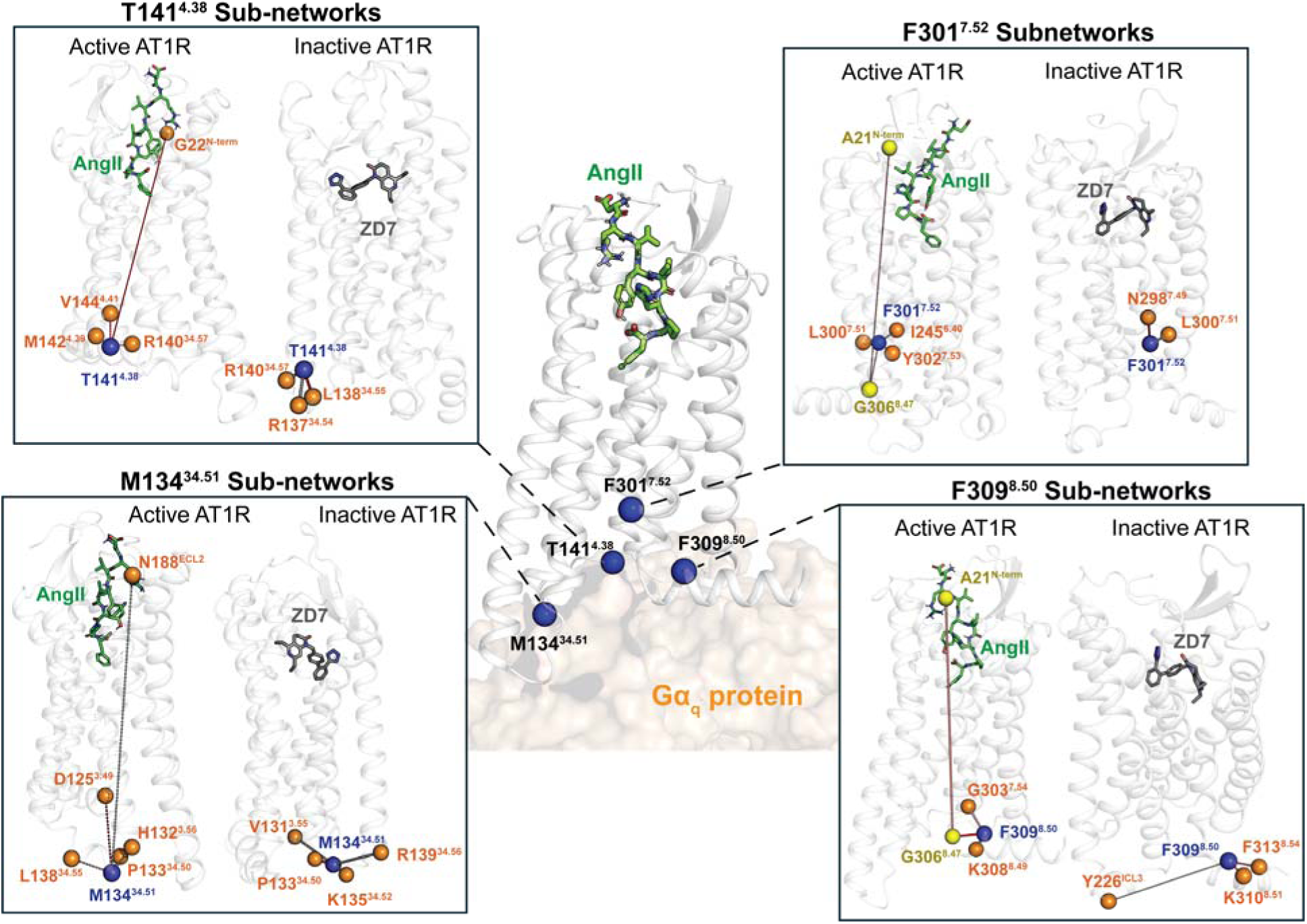
Bayesian network regions showing the allosteric effects of intracellular residues that modulate AngII binding. Four intracellular residues T141^4.38^, M134^34.51^, F301^7.52^, and F309^8.50^ affect AngII binding at a distance ^35^. Bayesian network analysis highlights sub-networks centered on these four residues that connect allosterically to the AngII binding pocket in the active state of AT1R. Mutagenesis studies show that perturbations at these sites alter AngII binding (Supplementary Fig. 10 in ^35^) and EC_50_ values, consistent with their predicted allosteric roles (see Fig. 3).

### Identification and characterization of a small molecule modulator, Q2 of AngII:AT1R

We next tested whether residue communities implicated in Gα_q_ coupling could be leveraged to find small molecule modulators of AngII:AT1R:Gα_q_ signaling. We clustered the active-state MD simulation trajectories of AT1R by root mean squared deviation (RMSD) in Cα atom coordinates and this resulted in 10 conformational clusters (details in Methods). We extracted a representative conformation for each of the ten conformational clusters and used the in-house software, *FindBindSite* ^39^ on each representative structure to identify putative small molecule binding sites (see Methods for details). The spatial locations of the diverse set of small molecule binding pockets found across the ensemble are shown in Fig. 6a. We prioritized pockets that were consistently present across highly occupied (>75% of the population) conformational clusters from the MD ensemble (Supplementary Fig. 7) and overlapped with the community of the top quartile of high positive-ΔWd residues identified by the DBNM (Fig. 6c). This overlap ensured that the selected pocket was not only persistent across the dynamic ensemble, but also functionally enriched for key residues involved in allosteric co-dependencies with other residues. Structure-based virtual screening of a 756,340 compound Enamine library against this pocket yielded 14 high-scoring candidates, selected based on docking rank, physicochemical filters, and overlap with DBNM-derived interaction hotspots (Fig. 6b). The rotamer conformation of the residues in the binding site of the selected hit molecules was optimized (see Methods for details). Nine commercially available compounds (Q1 to Q9; Supplementary Table 2) were evaluated for modulation of AT1R-mediated Gα_q_ activation using a Gα_q_/P63 translocation BRET biosensor ^40^. Preincubation of AT1R-expressing cells with 200 μM Q2 produced an approximately two-fold right-shift in the EC_50_ of AngII-induced Gα_q_ signaling (Fig. 6d, e). Although Q8 and Q9 increased maximal Gα_q_ responses by ∼25%, they were excluded from further study due to compromised luciferase activity and poor solubility (precipitation/aggregation) (Supplementary Table 2). Q2 was therefore selected for further characterization (Fig. 6f). The modest inhibitory potency of Q2 is consistent with it being a fragment with low molecular weight (296.32 Da), non-optimized scaffolds that typically exhibit weak binding affinity ^41^.

**Figure 6.**
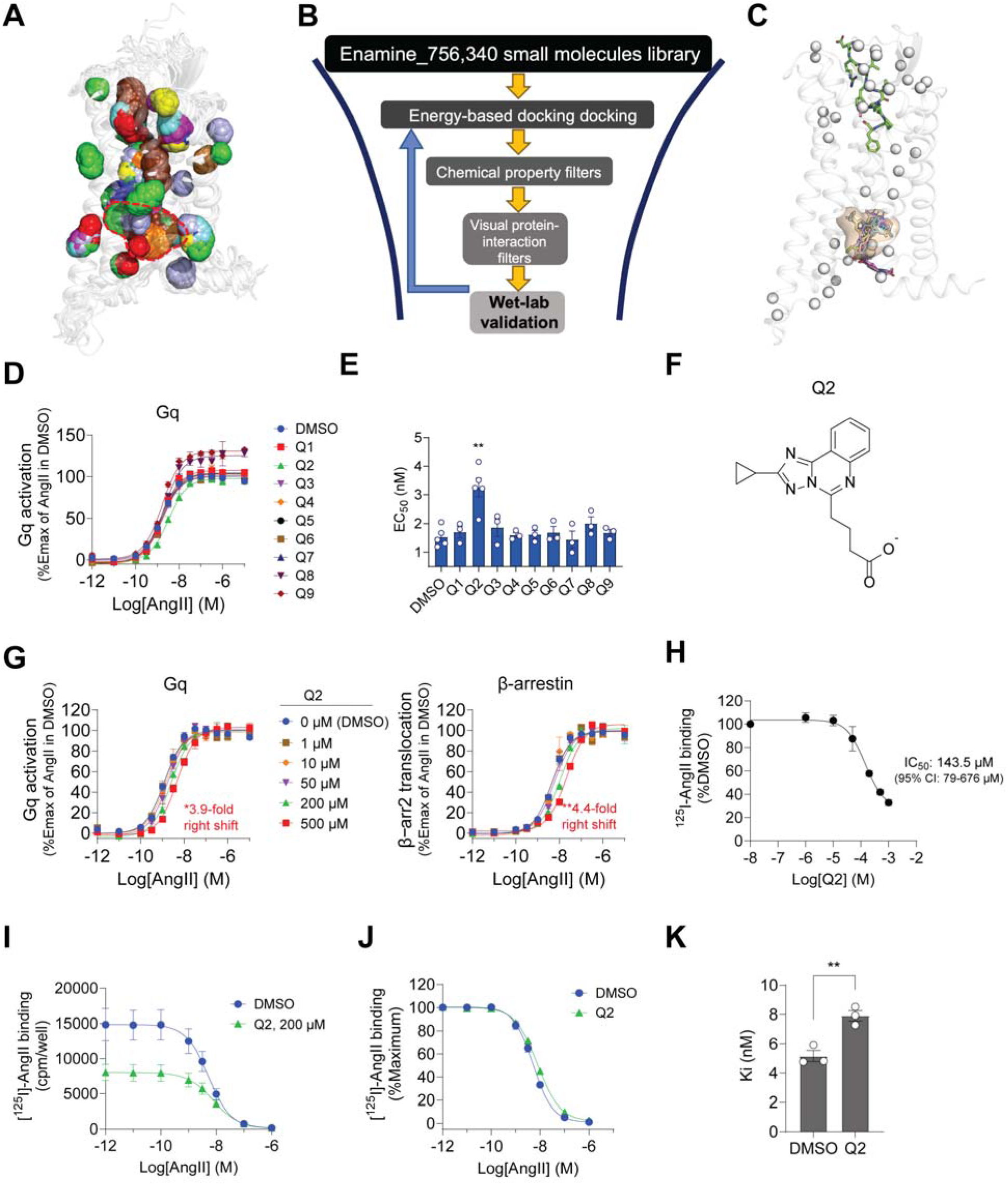
Identification of the allosteric pocket and allosteric modulator Q2 in AT1R. (**a**) Ten representative structures overlaid from the ten conformational clusters of the active state MD simulation ensemble of AngII and Gα_q_ bound AT1R used for identifying putative allosteric small molecule binding sites. The predicted putative small molecule binding pockets are shown as colored spheres. Each binding pocket is shown color-coded per conformation, with the merged overlay highlighting consensus binding pockets. The prioritized allosteric pocket (highlighted by red dashed line) was selected based on overlap with the residues in the top quartile of DWd from BN analysis. (**b**) Workflow for Virtual Ligand Screening (VLS) starting from ∼756,000 small molecules (details in the Methods section). Ligands were filtered through Glide docking, physicochemical property thresholds, pocket-specific filters, and validated against BN predictions and mutational data. (**c**) The selected allosteric pocket ‘Q’ shown in (a), together with the poses of top nine tested compounds. The top quartile DWd residues are shown in white spheres.(d) AT1R-mediated Gq activation as measured by Gq BRET assays. HEK293 cells expressing AT1R and the Gq BRET biosensor (Gα_q_, P63-RlucII, and rGFP-caax) were pretreated with 200 μM of Q compounds before AngII stimulation. Normalized BRET dose–response curves are shown relative to vehicle control. Data are from more than three independent experiments and expressed as mean ± SEM.(e) Plot of EC_50_ in nM from concentration response curves of AngII-mediated Gq activation. **, p < 0.001; unpaired Student’s t-test, compared to the value of DMSO. (**f**) 2D structure of Q2. (**g**) Concentration dependent effect of Q2. AT1R along with either Gq sensor (left) or b-arr2 sensor (right) transfected cells were preincubated with different concentrations of Q2 then stimulated with various concentrations of AngII. Data are mean ± SEM of three independent experiments. **(h**-**k**) Competitive displacement assay of radiolabeled AngII by Q2. (**h**) Competition binding of ^125^I-AngII to AT1R was performed in intact HEK293 cells in the presence of increasing concentrations of Q2. Data are expressed as percentage of maximal specific binding relative to vehicle (DMSO). Data represent means ± SEM of three independent experiments. (**i**) Competition binding of ^125^I-AngII to AT1R was performed in intact HEK293 cells using increasing concentrations of AngII in the absence or presence of 200 µM Q2. Data represent total binding (cpm/well) and are shown as mean ± SEM from three independent experiments. (**J**) Data in (I) were normalized as a percentage of its own maximum binding. (**k**) Estimated Ki values obtained from the curves in (J). Data are expressed as mean ± SEM with individual values. **, p <0.01, unpaired Student’s t-test, compared to the value of DMSO.

### The fragment Q2 acts a NAM to both G**α**_q_ and **β**-arr2 coupling to AT1R and its putative binding site is allosteric to AngII binding site

Q2 acts as a negative allosteric modulator (NAM) of AT1R by inhibiting Gα_q_ coupling through a predicted binding site topologically distinct from the AngII binding pocket. Dose dependent measurements of the effects of Q2 on AngII-induced Gα_q_ signaling showed ∼ 4-fold rightward shift in the AngII EC_50_ at the maximal soluble concentration (500 μM; Fig. 6g and Table 1). In radioligand binding assays, saturating concentrations of Q2 partially inhibited ^125^I-AngII binding to AT1R, with an apparent IC_50_ of ∼143 μM (Fig. 6h). Q2 alone reduced radioligand binding by ∼50% (Fig. 6i), yet only modestly altered AngII affinity in competitive assays (IC_50_ shift from 5 nM to 8 nM; p < 0.01; Fig. 6k). This limited displacement of AngII binding, together with the reduction in AngII potency, supports an allosteric mechanism of action. Q2 also reduced AngII-induced β-arrestin2 coupling to AT1R (Fig. 6g), indicating that modulation of this intracellular allosteric site impacts multiple downstream signaling pathways rather than selectively targeting Gα_q_, although with potential different mechanisms. However, in our previous study we have shown that the homology modeling of AT1R with β-arrestin2 does not recapitulate the alanine scanning BRET data for β-arrestin coupling ^35^ adequately and hence not suitable for performing BNM analysis for mechanistically understanding Q2’s mode(s) of action on this pathway. We therefore focused on understanding how Q2 affected AngII-AT1R-mediated Gα_q_ signaling.

**Table 1.** Q2 concentration dependent EC_50_ changes in AngII-mediated Gq and b-arrestin activation.

| Q2 concentration | Log$EC_{50}$ ($EC_{50}$ in nM) | | $EC_{50}$ Fold change | |
| --- | --- | --- | --- | --- |
|  | Gq | b-arrestin | Gq | b-arrestin |
| 0 mM (DMSO) | -8.92 ± 0.04<br>(1.2) | -8.32 ± 0.03<br>(4.8) | 1 | 1 |
| 1 μM | -8.93 ± 0.06<br>(1.2) | -8.34 ± 0.06<br>(4.6) | 1 | 1 |
| 10 μM | -8.85 ± 0.07<br>(1.4) | -8.36 ± 0.07<br>(4.4) | 1.2 | 0.9 |
| 50 μM | -8.76 ± 0.05<br>(1.7) | -8.24 ± 0.06<br>(5.7) | 1.4 | 1.2 |
| 200 μM | -8.61 ± 0.05**<br>(2.5) | -7.99 ± 0.05**<br>(10.3) | 2.0** | 2.2** |
| 500 μM | -8.33 ± 0.05***<br>(4.7) | -7.68 ± 0.04****<br>(21.1) | 3.9*** | 4.4**** |

To procure a validation of the predicted allosteric binding site of Q2 and NAM effect on Gq signaling, we first tested alanine substitutions of residues lining the docked Q2 pocket in the AngII-bound AT1R model (Fig. 7a, b). AT1R mutations produced different effects on Q2 mediated activity, consistent with an allosteric mechanism. Substitutions at F248A^6.43^, N295A^7.46^, and F301A^7.52^ abolished the NAM effect on AngII–Gα_q_ signaling (Fig. 7a, red sticks; Fig. 8a), indicating that these residues are critical for Q2 binding and/or affecting the NAM activity of Q2. In contrast, mutations at L119A^3.43^, R126A^3.50^, P299A^7.50^, and L305A^7.56^ enhanced Q2-induced rightward shifts in AngII potency, while substitutions at L70A^2.46^, N294A^7.45^, and N298A^7.49^ selectively decreased AngII efficacy (Emax) (Fig. 8 and Supplementary Table 3).

**Figure. 7.**
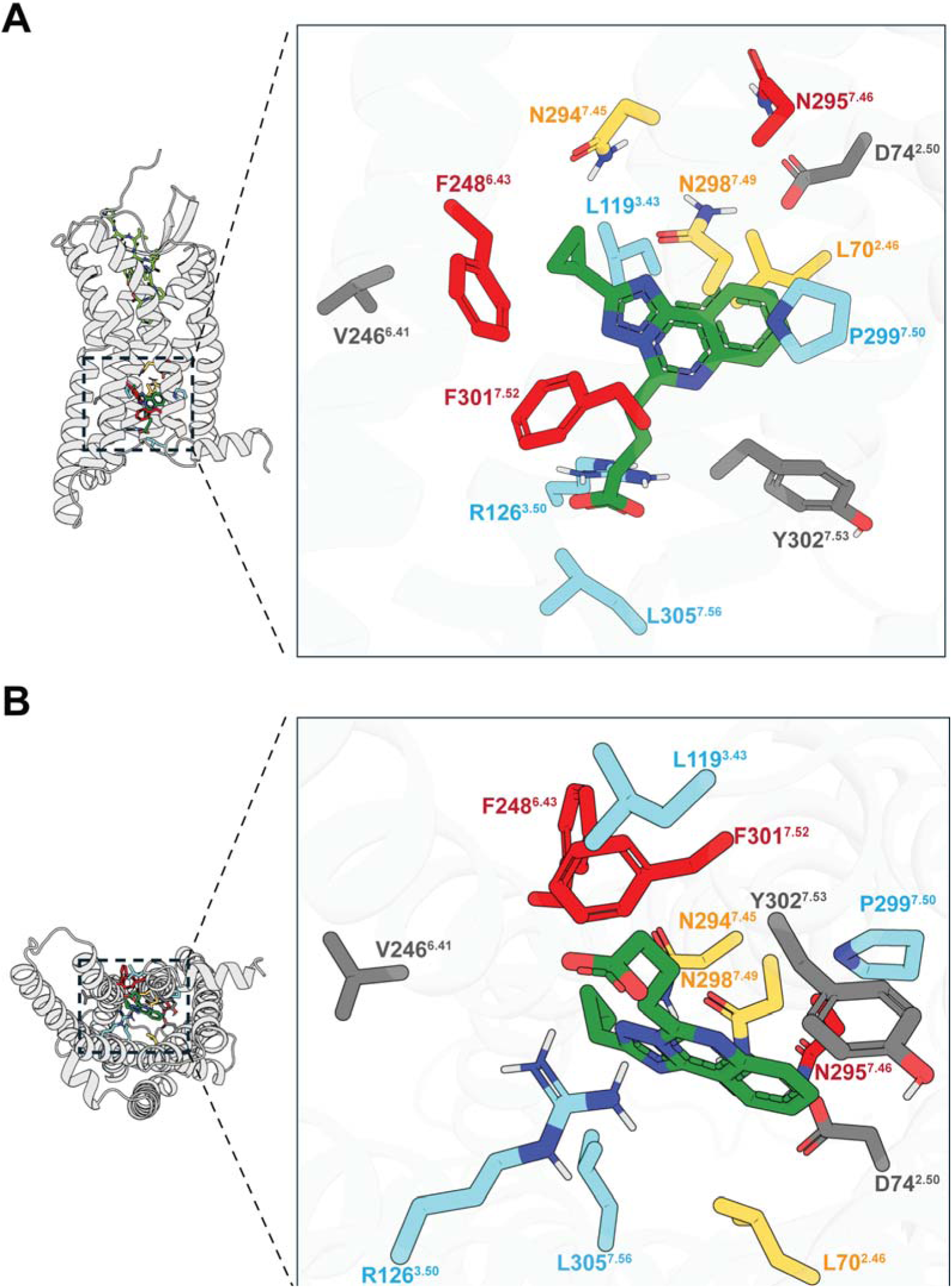
Characterization of the fragment Q2 allosteric binding pocket. (**a**) Front and (**b**) Bottom view of residues surrounding the predicted Q2 (green sticks) binding site in AT1R and tested experimentally. Residues labeled in red represent mutations that abolished Q2 NAM activity on AngII; residues in blue enhanced the Q2 NAM effect upon mutation; residues in yellow show changes in efficacy of Q2 mediated NAM effect and residues in gray indicate mutations without significant functional effect.

**Figure 8.**
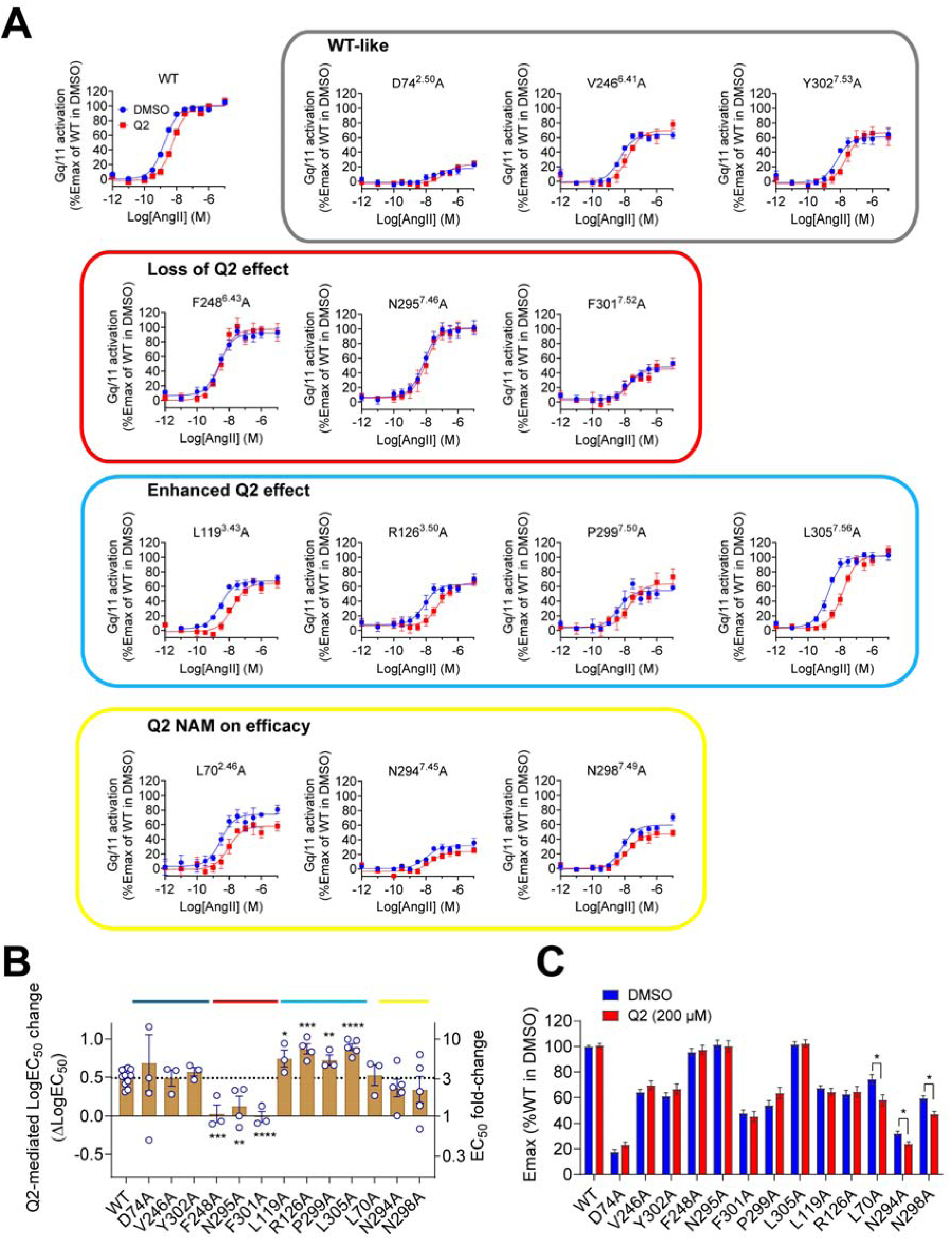
Experimental validation of the predicted allosteric binding site of Q2. (**a**) AngII-concentration response curves of the AT1R mutants. Data are normalized to the Emax of WT in vehicle control (DMSO) and expressed as mean ± SEM of more than three independent experiments. (**b**) Q2-mediated logEC_50_ changes. LogEC_50_ of each mutant’s AngII responses in the absence (DMSO) or presence of Q2 (200 μM) were obtained and calculated Q2-mediated logEC_50_ changes by subtracting logEC_50_ value of DMSO from that of Q2 treated. *, p < 0.05; **, p < 0.01; ***, p < 0.001; ****, p < 0.0001, unpaired Student’s t-test, compared to the value of WT. (**c**) Emax of AngII-mediated Gq/11 activation of the mutants in the absence or presence of Q2. The maximum asymptote of each curve in (A) is plotted. *, p < 0.05, unpaired Student’s t-test, compared to value of control (DMSO).

To next elucidate the mechanistic origins of the enhanced NAM effect of Q2 observed for the R126A^3.50^ and L305A^7.56^ mutants, we performed all-atom MD simulations of WT and mutant AT1R in complex with AngII, Gα_q_βγ, and Q2, followed by contact analysis between Q2 and its surrounding residues, and conformational analyses of individual side chain rotamers and α5 helix (Fig. 9 and Supplementary Fig. 9). In the active WT receptor, the intracellular TM5–TM7 interface is stabilized by a conserved packing interaction involving Y215^5.58^ and Y302^7.53^ that contributes to maintaining an active-like intracellular architecture. While this interaction is not captured as a persistent direct hydrogen bond, its stability is supported by a surrounding polar network in which R126^3.50^ plays a central organizing role. Positioned at the convergence of TM3, TM5, and TM7, R126^3.50^ can transiently engage multiple polar partners, enabling it to function as a relay that reinforces intracellular packing through ensemble-averaged interactions rather than a single static contact. Upon Q2 binding, R126^3.50^ forms a strong and persistent polar interaction with the small molecule, competitively sequestering its hydrogen-bonding capacity. This interaction rewiring weakens the native polar network that supports Y215^5.58^ – Y302^7.53^ packing, resulting in an outward displacement of TM7, pushing the receptor to a more inactive-like conformation (Supplementary Fig. 9A, B) and a remodeled intracellular cavity. In the WT receptor, Q2 remains anchored by R126^3.50^ (Fig. 9a, Supplementary Fig. 9C), restricting its depth of penetration toward the distal segments of the Gα_q_ α5 helix. Accordingly, principal component analysis (PCA) of α5 helix conformations reveals that the Q2-bound WT ensemble substantially overlaps with the WT active-state distribution, indicating preserved receptor–G protein alignment despite altered intracellular dynamics (Fig. 9c). Together, these observations support a model in which Q2 acts as a negative allosteric modulator by rewiring intracellular polar interaction networks rather than directly and sterically blocking G protein engagement.

**Figure 9.**
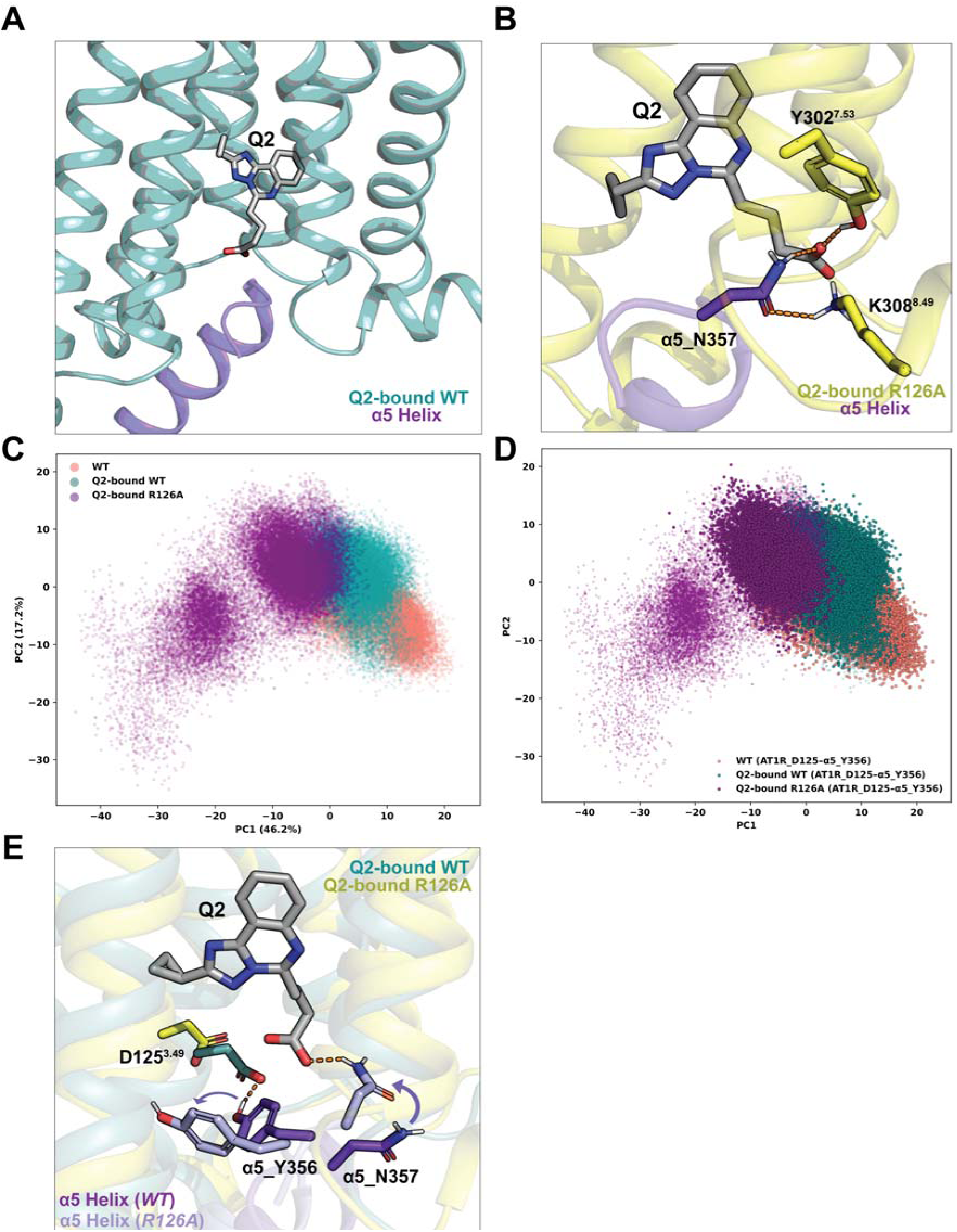
Q2 destabilizes productive G protein coupling by disrupting a conserved α5 polar lock revealed by the R126A mutant. (**a**) Structural view of Q2 bound to the intracellular allosteric pocket of AT1R in the WT receptor–G protein complex, shown relative to the Gα_q_ α5 helix. (**b**) In the R126A^3.50^ mutant, Q2 forms additional interactions with residues lining the intracellular pocket and engages the α5 helix through contacts involving N357, illustrating ligand proximity to the coupling interface. (**c**) Principal component analysis (PCA) of α5 helix conformations for WT, Q2-bound WT, and Q2-bound R126A systems. (**d**) Conditioning the α5 PCA on the presence of the conserved receptor–G protein D125– Y356 polar contact. (**e**) Structural comparison of representative WT-like and R126A α5 conformations highlighting the conserved D125^AT1R^–Y356^α5^ polar interaction that stabilizes productive coupling in the WT complex and is destabilized in the R126A mutant.

In contrast, R126A^3.50^ disrupts polar anchoring interaction that constrains Q2 in the WT receptor, resulting in increased positional and orientational freedom of the small molecule within the intracellular cavity of AT1R. In this context, Q2 more frequently adopts deeper poses toward the Gα_q_ interface and engages alternative polar contacts with residues such as Y302^7.53^ and K308^8.49^ (Fig. 9b, Supplementary Fig. 9F, G). Although Q2 - α5 hydrogen bonding via N357 is observed in this mutant, conditioning α5 helix conformational space on N357-Q2 hydrogen bond presence reveals that this interaction does not uniquely define the dominant off-pathway basin (Supplementary Fig. 9H). PCA demonstrates that the R126A^3.50^ system samples a distinct α5 helix conformational ensemble that is not accessed in either the WT or Q2-bound WT simulations (Fig. 9c). We observed that formation of a hydrogen bond between Q2 and N357 straightens the normally “wavy hook” region of the Gαq α5 helix, displacing it away from the GPCR binding cavity. Concomitantly, the stabilizing polar interaction between AT1R D125^3.49^ and Gα_q_ Y356 is disrupted (Fig. 9e). Mapping polar contacts between AT1R D125^3.49^ and Gα_q_ Y356 onto the α5 conformational landscape reveals a strong correspondence between loss of this receptor - G protein interaction and occupation of the R126A-specific basin (Fig. 9d and Supplementary Fig. 9I). Together, these results indicate that loss of the anchoring constraint in R126A^3.50^ permits Q2 to redistribute α5 helix conformational populations away from productive coupling geometries, thereby enhancing negative allosteric modulation through decoupling of receptor–G protein engagement rather than stabilization of a single inhibitory contact.

By contrast, the enhanced NAM activity of the L305A^7.56^ mutant reflects a distinct, network-level mechanism centered on how Q2 engages the conserved intracellular tyrosine module. In the WT receptor, R126^3.50^ likely contributes to organizing the local polar environment that supports the conserved Y–Y lock at the intracellular face. In the presence of Q2, R126^3.50^ is preferentially recruited into a persistent polar interaction with the ligand, biasing the intracellular bundle toward a Y–Y–disrupted arrangement rather than directly occluding the G protein interface. In L305A^7.56^, removal of the bulky leucine side chain induces a reorganization of the TM3/TM5/TM7 bundle (Supplementary Fig. 9E) that brings Y215^5.58^ into closer register with the Q2 pocket. As a result, Q2 forms a stabilizing hydrogen bond with Y215^5.58^ while retaining its interaction with R126^3.50^ (Supplementary Fig. 9D, E), effectively creating a R126–Q2–Y215 polar triad. This triad functionally sequesters Y215^5.58^ from coupling with Y302^7.53^, stabilizing the “unlocked” intracellular microstate and propagating conformational rearrangements across the TM5–TM7 interface. Thus, the stronger NAM phenotype in L305A^7.56^ is best explained not by a simple increase in small molecule’s affinity, but by reinforcement of a specific intracellular polar-network configuration that more strongly biases receptor dynamics away from productive coupling transitions.

Together, these results delineate two mechanistically distinct routes by which Q2 enhances NAM activity on AT1R signaling. In the R126A^3.50^ mutant, loss of the native polar hub redistributes Q2 within the intracellular cavity, leading to conformational decoupling of the Gα_q_ α5 helix through disruption of productive receptor–G protein contacts. In contrast, the enhanced NAM activity observed in L305A^7.56^ arises from mutation-induced rewiring of the intracellular allosteric pocket, whereby Q2 stabilizes a specific polar network that biases the receptor toward an intracellularly “unlocked” microstate without directly blocking G protein engagement. Consistent with this framework, mutations that abolish NAM identify residues that contribute directly to Q2 recognition within the pocket, whereas mutations that enhance NAM reveal residues that normally restrain allosteric coupling in the WT receptor. Collectively, these structural and functional studies show that Q2 NAM activity is modulated by the mutations of the residues in the binding site thereby validating the predicted binding site and the allosteric binding mode of action of Q2 on AT1R:G_q_ signaling. This also highlights how our integrated MD–DBNM approach can uncover cryptic allosteric sites and provide a framework for rational modulation of GPCR signaling.

## Discussion

AT1R signaling arises from ensembles of interconverting conformations ^38,42–44^. Yet the field has lacked robust methods capable of resolving how information is routed through this dynamic landscape to control G-protein coupling ^37,45^, which can be harnessed to identify cryptic and allosterically communicating modulator binding sites to control receptor signaling. By integrating Differential Bayesian Network Model (DBNM) applied to MD ensembles with whole-receptor mutational functional scanning, and through iterative prediction and experimental testing of residues that modulate Gα_q_ coupling using deep mutational scan, we generated a residue-level map of the conditional co-dependencies in interaction energies that underlie the allosteric communication. This framework moves beyond correlation-based descriptors and static structural comparisons to identify residues whose influence emerges only through their participation in state-dependent information flow.

Consistent with this network-centric view, we show that mutations in flexible or intrinsically disordered regions of AT1R produced large effects on Gα_q_ coupling, revealing a role of these dynamic and understudied segments in modulating AT1R Gα_q_ coupling. These results establish an allosteric mechanism of AT1R activation and enable us to uncover an intracellular allosteric site that can be targeted by a small molecule, whose functional role had remained unresolved despite extensive structural characterization.

The BNM on active and inactive states of AT1R produces a sparse, causally grounded graph on each state that assigns each residue a probabilistic “influence” score (Wd) while filtering out indirect or spurious edges. Using this framework, we show that AT1R distributes information flow across mediator communities bridging the ligand binding site (LBS) and G protein interface (GPI). The residues involved in the allosteric communication do not necessarily contact AngII or Gα_q_ directly, yet they occupy high co-dependent positions in the network that distinguish the active and inactive ensembles. Importantly, because the BNM aims to capture only direct (non-transitive) dependencies, it explains why certain alanine substitutions produced no changes in Gα_q_ coupling, whereas other positions appear insensitive only because alanine is a conservative mutation. Testing this buffering effects with predicted non-alanine mutations showed distinct effects on Gα_q_ coupling. This ability to distinguish between (i) true insensitivity, (ii) network buffering effects, and (iii) direct co-dependencies enables us to readily translate mutational data into mechanistic hypotheses about the structural logic of AT1R signaling.

Furthermore, we show that residue communities that play a dominant role in allosteric communication in the active state and coincide with transient, cryptic pockets are potential modulator sites. These pockets sit at communication bottlenecks where a small molecule could shift signaling by perturbing information flow rather than by obstructing AngII binding. This mechanistic framework explains how mutations far from the orthosteric pocket alter Gα_q_ coupling/ligand binding, and it highlights structurally connected, dynamically encoded pockets as promising domains for allosteric modulation targeting with drugs.

Most structure-based network analyses, including contact maps, co-fluctuation networks, and recent difference contact network analyses ^46–48^, identify where conformational changes occur but cannot distinguish direct from indirect associations. As a result, they often inflate correlations driven by multi-collinearity within the receptor core. By inferring conditional rather than correlative dependencies, the BNM resolves this ambiguity and identifies residues whose influence genuinely changes between states. This explains why ΔWd predicted both the established functional sites and the buffered, alanine-insensitive positions later validated by non-alanine mutational testing.

The DBNM’s predictions held up prospectively. Residues with the strongest ΔWd values were frequently the same positions whose alanine substitution impaired Gα_q_ coupling, and quartile clustering analyses showed that top-ranked nodes aggregate in the LBS, mediator, GPI, and sodium-site neighborhoods, regions expected from an allostery model ^45,49^. Equally important, the DBNM pinpointed a subset of high ΔWd residues with no change in G protein coupling with alanine substitution, yet whose targeted non-alanine substitutions produced clear Gα_q_ coupling changes. These results reveal that sidechain and not merely backbone presence, governs modulation of activity at these residue positions. Thus, the BNM complements alanine scanning by rescuing functional false negatives and enabling a more complete, mechanistically insightful map of functionally critical residues.

Functional mapping of key residue communities involved in Gα_q_ coupling enabled discovery of allosteric sites. Across active-state MD ensembles, ΔWd-enriched mediator clusters consistently overlapped with a small set of transient cavities adjacent to the GPI. Using this map as a guide, we identified Q2, a fragment-sized small molecule that showed attenuation of AngII-driven Gα_q_ coupling. Mutational analysis of residues lining the predicted cavity provided convergent evidence for a localized allosteric site: substitutions at several positions abolished or attenuated Q2’s NAM effect. Together, these data validate that the BNM-derived mediator pocket constitutes a bona fide allosteric site capable of modulating AT1R signaling.

Together, our DBNM framework extends beyond AT1R, enabling the identification of functionally critical residues with nonpenetrant mutational effects and the discovery of allosteric binding sites across GPCRs. By combining MD ensembles, mechanistically interpretable BNM inference, and targeted mutagenesis, it generates specific, testable, and mechanistically grounded hypotheses, providing a practical route for the rational design of GPCR allosteric modulators.

## Materials and Methods

### Computational methods and data analysis

#### MD simulations

MD simulations were performed using the GROMACS package (version 2016/2019) with the Chemistry HARvard Molecular Mechanics (CHARMM) 36 force field for proteins, palmitoyl-oleoyl-phosphatidylcholine (POPC) lipids, ions, and using CHARMM Transferable Intermolecular Potential with 3 Points (TIP3P) water as solvent ^50,51^. We performed separate MD simulations starting from the active-state cryo-EM structure of the AT1R-AngII-Gα_q_-Gβ_1_-Gγ_2_ complex (PDB ID: 7F6G), the inactive-state crystal structure of AT1R-ZD7 complex (PDB ID: 4YAY), the AT1R-AngII-Q2-G _q_-Gβ_1_-Gγ_2_ complex, the R126A_AT1R-AngII-Q2-Gα_q_-Gβ_1_-Gγ_2_ complex, and the L305A_AT1R-AngII-Q2-Gα_q_-Gβ_1_-Gγ_2_ complex. Sar^1^-AngII was reverted to AngII in 7F6G, and molecules other than listed proteins and ligands were removed in structure preparation. The soluble cytochrome b562 was removed in 4YAY, and ICL3 was reconstructed during structure preparation. The AT1R-Q2 complex was taken from the glide-dock results, with Gα_q_-Gβ_1_-Gγ_2_ complex remains the same as active-state structural setup. For AT1R mutants, the mutated residue is computationally swapped to Ala, and the side chain rotamer conformation of residues within 5Å of the swapped residue is energetically optimized. The hydrogen atoms were added, protein chain termini were capped with neutral acetyl and methyl amide groups, and histidine-protonated states were assigned using Maestro (Schrödinger). The simulation box was created using CHARMM-GUI. We used the OPM (orientation of proteins in membranes) structure of PDB: 7F6G for alignment of the TM helices of protein structure and inserted a pre-equilibrated POPC bilayer. Final system dimensions were as follows: AT1R-AngII-Gα_q_-Gβ_1_-Gγ_2_ 120 × 120 × 179 A□, including 371 lipids, 58170 waters and 150 mM NaCl; AT1R-ZD7 75 × 75 × 122 A□, including 130 lipids, 13801waters and 150mM NaCl; AT1R-AngII-Q2-Gα_q_-Gβ_1_-Gγ_2_ 135 × 135 × 185 A□, including 489 lipids, 78019 waters and 150 mM NaCl; L305A_AT1R-AngII-Q2-G□_q_-Gβ_1_-Gγ_2_ 140 × 140 × 186 A, including 531 lipids, 84956 waters and 150 mM NaCl; R126A_AT1R-AngII-Q2-Gα_q_-Gβ_1_-Gγ_2_ 140 × 140 × 186 A, including 533 lipids, 85312 waters and 150 mM NaCl. After minimizing the AT1R-ligand complex, we equilibrated it using an NVT (constant temperature, constant volume) ensemble (1 ns long) and consequently with an NPT (constant temperature, constant pressure) ensemble where we gradually reduced the position restraints from 5 to 0 kcal/mol per square angstrom (each step was 5 ns long). In the last step of equilibration, we performed 5 ns of unrestrained NPT simulations before running a total of five production MD simulations, each 1000 ns long. The snapshots were stored every 20 ps, and every 10^th^ snapshot of the entire 1000 ns × 5 runs amounting to 5 μs of simulation time was used for analysis (25,000 snapshots total).

#### Residue-wise interaction energy calculation

To quantify residue-level energetic contributions to allosteric communication, nonbonded interaction energies were computed for each AT1R residue across the MD trajectories. The nonbonded interaction energy (Eij) between any two residues (i and j) is composed of two separate energy terms: The van der Waals (vdW) interaction energy (VLJ) given by the Lennard-Jones (LJ) potential (Eq. 1) and the electrostatic interaction energy (VC) given by Coulombic potential (Eq. 2). Specifically, the short-range (within 12 Å) Coulombic and van der Waals forces were calculated using gmx energy module in GROMACS ^52^ and extracted from the energy log file. The computed interaction energy is defined as the sum of the LJ and Coulombic interaction energies (Eq. 3) for every MD snapshot (25,000). All favorable interaction energies are <0 kJ/mol and hence have a negative sign.

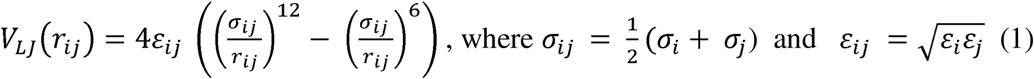

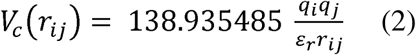

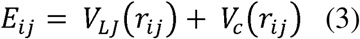

This procedure yielded a time series of interaction energy values for each residue, forming the input features for Bayesian network modeling. Distributions of these interaction energies for intracellular loop (ICL) residues and transmembrane (TM) residues are shown in Supplementary Fig. 1A, B.

#### Construction of differential Bayesian network models

DBNM starts by constructing two distinct *G_A_,G_B_* BN models for the two systems (datasets) to be contrasted. In this study, we have applied it to delineating the differences between the active state and inactive state of AT1R, aiming to identify the structural differences in the interaction networks. For each *G* we obtain the weighted degree for all nodes as 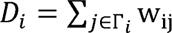, where *r_i_* is the set of neighbors of the node *i* and *w_ij_* is the commensurate, universal-scale, edge strength ^28^. We have previously demonstrated that the weighted degree *D_i_* is a strong correlate of a node’s functional significance, with high weighted degree predictive of nodes (both residues and residue contacts) playing key roles in protein function and stability ^53^ Here, we are interested in identifying salient nodes that also differentiate between the two systems being contrasted. To that effect, we define 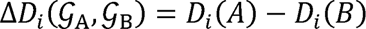 as the weighted degree difference for the node i between the two BN *G_A_*, *G_B_* This measure quantifies the extent of changes in the dependency structure as identified by BNs in the two systems, directly pointing to the most important differential modulator(s) for these two systems. Then, we identify the set vertex V*predominantly important for the system A (or B) as 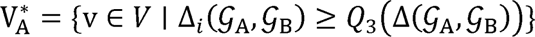, (and 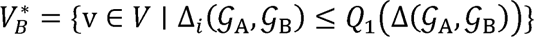 where *Q*_1_, *Q*_3_ are the first and third quartiles, respectively. The overlay of the V* nodes onto the 3D structures for the two systems allows visual identification of the protein substructures corresponding to the key differences.

### Testing robustness of BNM: bootstrap resampling of weighted degree and differential weighted degree

To evaluate the robustness of BN-derived residue influence, we performed a nonparametric bootstrap procedure in which the MD-derived residue interaction energy matrix was resampled 1,000 times. For each bootstrap replicate, 5,000 MD frames were randomly drawn with replacement from the 25,000-frame original trajectory. Pairwise mutual information (MI) between residues was recomputed for each resampled dataset. This procedure retains the dependency graph structure inferred from the full dataset while allowing edge weights to fluctuate according to sampling variability in the underlying MD ensemble. For each residue *r*, the 1,000 bootstrap replicate *i* values formed the distribution: 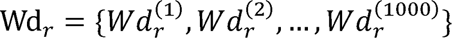 Mean Wd values were computed for active and inactive networks separately: 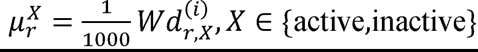. State-dependent influence shifts were quantified as: 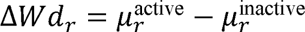. Bootstrap variances were propagated assuming independence of active and inactive estimates: 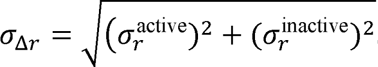. These propagated uncertainties were used to generate confidence intervals and assess the stability of state-dependent effects.

#### Structural partitioning and definition of allosteric dependencies

To obtain the list of amino acid residues that make contact between ligand-receptor and receptor-G protein we utilized the Get_Contacts software (https://getcontacts.github.io/). To interpret information flow across AT1R, residues were categorized into three structural regions: 1. Ligand-binding site (LBS): residues contacting AngII (active state) or ZD7155 (5,7-diethyl-1-[[4-[2-(2H-tetrazol-5-yl)phenyl]phenyl]methyl]-3,4-dihydro-1,6-naphthyridin-2-one) (inactive state) with greater than 20% frequency across MD snapshots. 2. G protein interface (GPI): residues contacting the Gαq subunit with >20% frequency in the active-state simulations. 3. Mediator residues: all remaining AT1R residues. Edges connecting residue pairs whose Cα atoms were separated by more than 10 Å were classified as long-range or “allosteric” dependencies. Edge strengths within and between these regions were summed to quantify patterns of information flow across receptor states.

#### Mutual Information (MI) analysis of residue pairs

To evaluate residue dependencies identified by the BN, we performed MI analysis on residue interaction energies extracted from MD simulations. MI was calculated for all residue pairs to capture statistical dependencies irrespective of their physical proximity. BN edge mapping was then combined with MI values to assess whether BN-highlighted interactions were supported by statistically significant dynamical coupling (Supplementary Fig. 8). Residue pairs with Cα–Cα distances greater than 30 Å were specifically examined to evaluate long-range communication (Supplementary Fig. 3). Structural mapping of the top 25% MI-supported, BN-derived residue pairs was conducted to visualize connections across functional regions, including ligand-binding sites, mediator regions, and G-protein interaction interfaces, in both active and inactive states (Fig. 2).

### RMSD-based clustering of the transmembrane backbone in active-state AT1R

RMSD-based clustering was performed using GROMACS rms module by aligning using backbone atoms of transmembrane residues (GPCRdb) and then clustering using a cutoff of 0.061 nm. The representative structure from each of the ten clusters were outputted as a PDB and rendered in Schrodinger PYMOL.

### Method of identifying druggable binding sites using the code FindBindSite

FindBindSite^39^ was used to identify putative druggable small-molecule binding pockets where potential allosteric modulators may bind. FindBindSite is a workflow that involves docking a small (60,000) and diverse library of small molecules to the entire protein structure. We used Glide to dock the 60,000-molecule library to AT1R. After docking this diverse library of 60,000 molecules, we clustered the regions with the highest docked ligand density and favorable dock score and ranked the sites. FindBindSite then outputs the rank of each site and the centers of the clusters that show ligand occupancy above the cut-off ligand density.

#### In silico deep mutational scanning

Computational mutational scanning was performed using the Schrodinger Maestro *Residue Scanning* module. Active-state crystal structure of the AT1R-AngII (PDB ID: 7F6G stripped of G _q_-Gβ_1_-Gγ_2_ complex) was loaded into the panel, and residues P19, K20, A21, H24, S107, V169, F171, Q187, N188, G194, L195, T198, P233, D236, D237, K240, were selected to be mutated to all other 19 amino acids. For each residue, all 19 possible non-native amino acid substitutions were modeled. For each mutant, two energetic metrics were computed: (i) the change in interaction energy of AT1R with Gα_q_ (ΔΔG_affinity_), and (ii) the change in total receptor stability of the mutant with respect to WT (ΔΔG_stability_). These values were summarized for the subset of mutants tested experimentally in Fig.4 and Supplementary Table 1. Postmutational rotamer conformation refinement was limited to within 5 Å of mutated residue. Stability was optimized with exhaustive enumeration. Candidate mutations were prioritized for experimental testing based on three criteria: (i) a significant change in AT1R-Gαq interaction energy (either stabilizing or destabilizing) relative to other substitutions at the same position; (ii) predicted receptor stability within 200 kcal/mol of the wild-type ensemble; and (iii) maintenance or strengthening of interhelical hydrogen bonds and van der Waals packing interactions relative to the wild-type structure.

#### Enrichment analysis

Residues were ranked by their differential weighted degree (ΔWD) values, calculated as the difference between the weighted degree in the active-state BN model and the inactive-state BN model. Experimental “positives” were defined as residues showing a statistically significant change (**p** < 0.05, Student’s *t*-test) in at least one pharmacological parameter (EC□□, Emax, or relative activity) compared to wild-type in the alanine scanning mutagenesis assays. For each integer *n* from 1 to the total number of ranked residues, the enrichment score (ES) was computed As 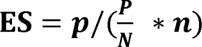,where n is sample size of the subset; N is total sample size; p is true positive rate of the subset; is total true positive rate. An ES > 1 indicates that the top-n residues are enriched for experimental positives relative to the background distribution. Enrichment curves were plotted by sequentially calculating ES for increasing n values, with every sixth point shown for visualization in Fig. 1.

#### Fisher’s exact test

To assess the statistical significance of enrichment at specific ranked subsets, Fisher’s exact test was performed comparing the number of experimental positives and negatives in the subset versus the remainder of the ranked list. The 2 × 2 contingency table consisted of: (i) Experimental positives in subset (ii) Experimental negatives in subset (iii) Experimental positives in remainder (iv) Experimental negatives in remainder. A one-tailed Fisher’s exact test (p < 0.05) was used to determine whether the proportion of positives in the subset was greater than expected by chance.

### MI and BNM analyses uncover allosteric communication pathways

Given the clear hierarchical clustering and differential connectivity patterns identified among residues in active versus inactive states, we next sought to quantitatively validate these interactions at the residue-pair level using MI analysis combined with BN edge mapping. MI captures statistical dependencies between residue pairs irrespective of their structural proximity, providing an additional validation layer for the BN-derived residue dependencies and their roles in state-specific receptor dynamics. Analysis of residue pairs highlighted by BN edges at long distances (>30 Å) revealed critical insights into genuine allosteric communication within AT1R. These residue pairs, despite their significant physical separation, are identified by our model to have dynamical dependencies, indicating that their interaction energies were statistically coupled in a state-dependent manner. This long-range coupling suggests these interactions facilitate communication between spatially distant functional regions of the receptor, aligning well with classical definitions of allosteric interactions. Structural mapping of these long-range interactions further supported their functional relevance, linking residues in ligand-binding and G-protein coupling sites through intermediary mediators in both active and inactive states (Supplementary Fig. 4).

### Experimental methods

#### Reagents

Human AngII [Asp^1^-Arg^2^-Val^3^-Tyr^4^-Ile^5^-His^6^-Pro^7^-Phe^8^], poly-L-ornithine, ANTI-FLAG® M2-Peroxidase (HRP) antibody, and SIGMAFAST OPD were purchased from Sigma-Aldrich. Dulbecco’s Modified Eagle Media (DMEM), fetal bovine serum (FBS), and other cell culture additives were purchased from Gibco, Life Technologies. Linear polyethyleneimine MW 25000 (PEI) was purchased from Polysciences. Sixteen percent paraformaldehyde (PFA) was purchased from ThermoFisher Scientific. BSA (Bovine Serum Albumin) was purchased from Wisent. Coelenterazine 400a was purchased from Nanolight Technology. Q5 high-fidelity DNA polymerase, Gibson Assembly Master mix, DpnI, and other PCR reagents were purchased from New England BioLabs.

#### DNA constructs and mutagenesis

Alanine mutants of Signal peptide-FLAG-tagged human AT1R (sp-flag-AT1R) construct were described previously ^35^. Gα_q_ DNA was obtained from cDNA.org (www.cDNA.org). P63-RlucII, βarr2-RlucII, and rGFP-CAAX were described previously ^40,54^. Mutations were introduced into the sp-flag-AT1R using a two-fragment PCR approach, as previously described ^55^. In short, forward and reverse site-directed mutagenesis primers (mutant-F and mutant-R) were designed with 21 bp of Gibson homology for Gibson assembly recombination. Stepdown PCR was used to make mutations, where two separate PCR reactions (primer pairs of mutant-F with Ori-R primers and mutant-R with Ori-F primers, respectively) were carried out to divide the vector in half. Each half of the PCR samples thereafter was purified and Gibson assembled. Reassembled vectors were transformed into *E. coli* and colonies were selected and amplified. Mutations were confirmed by Sanger sequencing (Genome Quebec CES). All primer sequences are listed in the Data S3.

#### Cell culture and transfection

HEK293 cells were cultured in DMEM supplemented with 10% FBS and 20 μg/ml gentamicin. Cells were authenticated (GeneCopoeia™) and tested for mycoplasma contamination periodically (ABM™, Mycoplasma PCR Detection kit). Cells were grown in 5% CO_2_ and 90% humidity. Cells were seeded at a density of 9,000 cells per well in a white (for BRET) or clear (for ELISA) 96-well plate, and the next day, transfected with receptor and BRET sensor constructs using PEI transfection reagent. Briefly, 1 μg of total DNA in 100 μl of PBS was mixed with 100 μl of PBS containing 2.5-3 μl of 1 mg/ml PEI. For Gq signaling (Gα_q_/P63 sensor), 100 ng of AT1R DNA with 20 ng of Gα_q_, 30 ng of P63-RlucII, and 80 ng of rGFP-CAAX were used. For the β-arr2 sensor, 150 ng of AT1R, 20 ng of βarr2-RlucII, and 80 ng of rGFP-CAAX DNA were used. For Gq/11 signaling by endogenous Gα_q_ and Gα_11_, 150 ng of AT1R DNA with 30 ng of P63-RlucII and 80 ng of rGFP-CAAX were used. For ELISA, 300 ng of AT1R DNA was used. Empty pcDNA was used to make up 1 µg of the total DNA amount. After 20 min incubation, the DNA/PEI complexes were dispensed into cells in 96-well plates (15 µl/well). All assays were performed 48 h post-transfection.

#### BRET measurements

HEK293 cells transfected in a white 96-well plate were washed once with Tyrode’s buffer (140 mM NaCl, 2.7 mM KCl, 1 mM CaCl_2_, 12 mM NaHCO_3_, 5.6 mM D-glucose, 0.5 mM MgCl_2_, 0.37 mM NaH_2_PO_4_, 25 mM HEPES, pH 7.4) and left in Tyrode’s buffer. Coelenterazine 400a (final concentrations of 2.5 µM) was added 1 min before ligand stimulation. Cells were then stimulated with various concentrations of AngII for 3 min for the Gα_q_/P63 sensor or 5 min for the b-arrestin translocation sensor, prior to BRET measurements. For compound treatment, compounds were preincubated at 37 °C for 20 min, then 10 more min at RT before ligand stimulation. BRET signals were measured using a Synergy2 (BioTek) microplate reader with emission filters, 410/80 nm and 515/30 nm for detecting the RlucII (*Renilla* luciferase) (donor) and GFP10/rGFP (acceptor) light emissions, respectively. The BRET ratio was determined by calculating the ratio of the light emitted by GFP10/rGFP over the light emitted by the RlucII. The effects of the compounds on luciferase activity were quantified based on the 410-nm readout.

#### Whole-cell ELISA

HEK293 cells were transfected with pcDNA, WT, or mutant receptors in poly-L-ornithine-coated, transparent 96-well plates to quantify the expression of receptors on the cell surface. 48 hours after transfection, cells were washed with PBS containing 1 mM MgCl_2_ and 1 mM CaCl_2_ (PBS-CM) and fixed with 4% PFA. Following blocking with 1% BSA in PBS-CM, cells were treated with ANTI-FLAG® M2-HRP antibody (1:4000) for one hour. After two PBS-CM and two PBS washes, 100 µl of SIGMAFAST OPD solution was added to each well. In order to stop the reaction, after 10 min, 25 μl of 3 M HCl was added. Then a Synergy 2 microplate reader (Bio-Tek) was used to read the plate at an absorbance of 492 nm. The non-specific signal from pcDNA-transfected cells was deducted to acquire a specific signal.

#### Intact cell radioligand binding

[^125^I]-AngII was prepared with the iodogen method as previously described (*29*). AngII binding and dose displacements were done using [^125^I]-AngII as a tracer. HEK293 cells were seeded at a density of 1 × 10^6^ cells per 100 mm dish, and the next day, cells were transfected with 3 mg of AT1R along with 3 mg of empty pcDNA using PEI methods (DNA: PEI, 1:2 ratio). The following day, cells were detached using TrypLE and re-seeded onto poly-L-ornithine-coated 24-well plate at a density of 1.5-2×10^5^ cells per well. The next day, cells in a 24-well plate were washed once with PBS, then incubated in binding buffer (50 mM Tris, 100 mM NaCl, 5 mM MgCl_2_, 0.2% BSA, pH 7.2). Dose displacement experiments were done by incubating the cells for 1 h at room temperature with ∼ 0.1 nM [^125^I]-AngII as tracer and increasing concentrations of Q2 or AngII in the absence (DMSO) or presence of 200 mM of Q2. Q2 was preincubated at 37□°C for 5–10 minutes, followed by cooling to room temperature for 10 min before the addition of AngII or [¹²□I]-AngII. After a 1-hour incubation period, the cells were washed three times with ice-cold PBS to remove any unbound radioactivity. Cells were solubilized in 0.5□M NaOH/0.05% SDS, and radioactivity was counted using a Wizard 1470 automatic γ-counter.

#### Data analysis

Data were normalized to the maximal response (Emax) of AngII for the WT receptor or WT receptor in control (vehicle only) treatment. LogEC_50_ and Emax were derived from BRET concentration-response curves using GraphPad Prism 10 software (GraphPad Software Inc.). Curves represent the best fits. The data from a typical BRET experiment followed a normal probability distribution, and three experiments were considered sufficient to accurately estimate the SD. Student’s *t* tests were therefore used to compare differences in the logEC_50_, Emax, and log (Emax/EC_50_) of each mutant to WT in the same pathway, and RAs.

## Acknowledgments

We thank Dr. Supriyo Bhattacharya for his valuable input on this work. This work was funded by grants from the National Institutes of Health grants R01-GM117923, R35 GM156498 to N.V., NLM R01LM013138 grant to A.S.R., and NIH NLM R01-LM013876 to N.V., A.S.R and S.B., Dr. Susumu Ohno Chair in Theoretical Biology (held by A.S.R.), Canadian Institutes of Health Research (PJT-162368 and PJT-173504 to S.A.L.)

## Author contributions

HC developed the DBNM framework, and performed computational analyses including ΔWd hotspot identification, virtual ligand screening for candidate allosteric modulators, MD simulations of mutant AT1R systems to elucidate the mechanism of Q2 modulation. YN performed experimental evaluation of small-molecule effects on WT and mutant AT1R signaling. ZAJ experimentally tested receptor mutations predicted from the DBNM hotspot analysis. WJCvdV performed the WT AT1R MD simulations. EM contributed to early methodological development. GK developed robustness test for the networks. HC, YN and ZAJ prepared the figures and wrote the manuscript with input from all authors. HC, YN, EM, ASR, SB, SAL and NV conceptualized the study. ASR, SB, SAL, and NV supervised the project and secured funding. All authors discussed the results and approved the final manuscript.

## Data and materials availability

The code for running the BN models is available in GitHub: https://github.com/bandyt-group/bandyt. Networks for active- and inactive-state AT1R provided as. graphml files as supplementary material. Experimental data have been provided to the journal as attachments.

## Supplementary Materials

In a separate file

