## Supplemental figures and tables for "Interpretable Machine Learning Model of Receptor Dynamics Reveals AT1R Allostery and a Negative Allosteric Modulator"

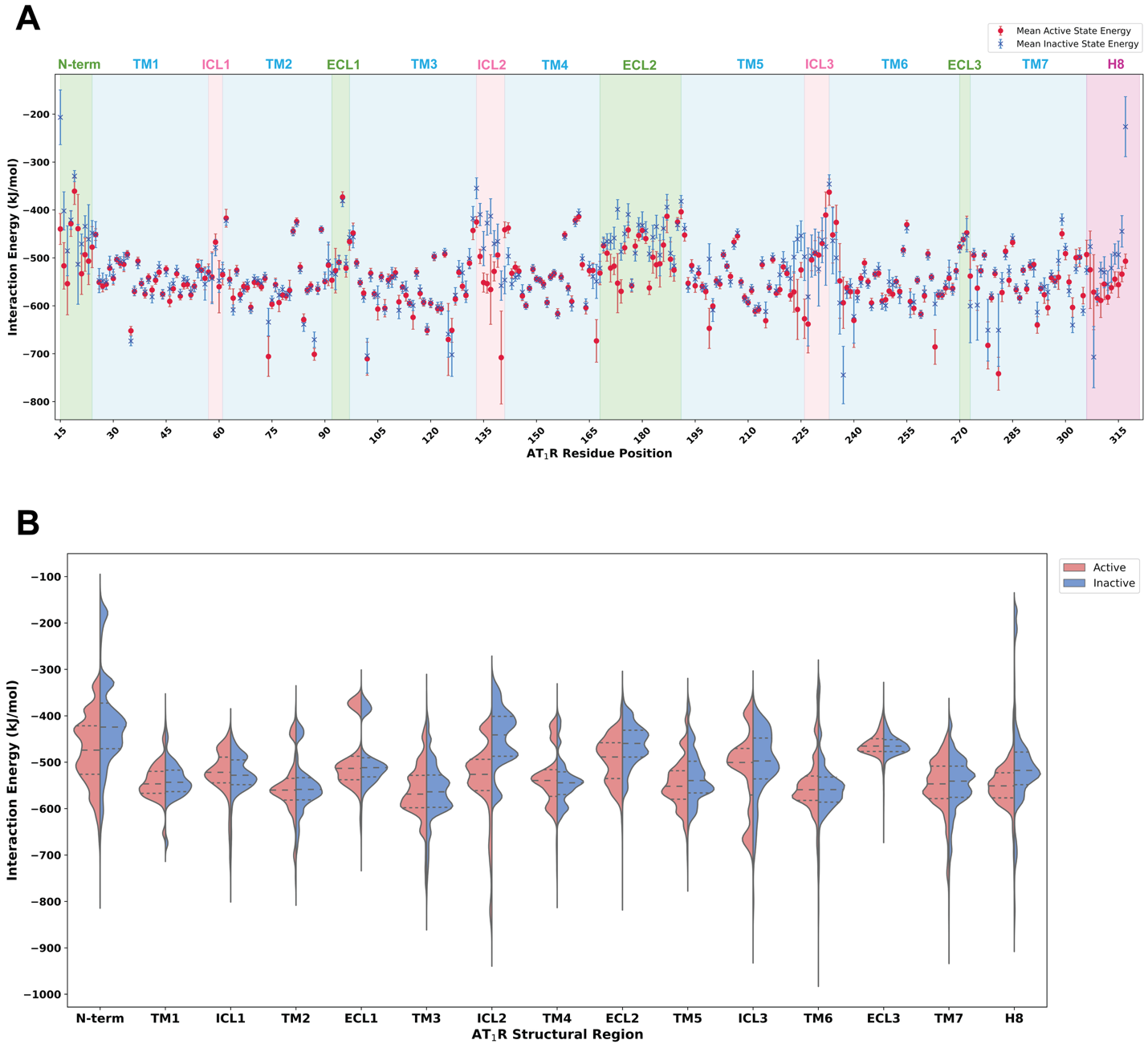

**Supplementary Fig. 1.** **Detailed residue interaction energy profiles extracted from MD simulations of active and inactive AT1R states.** (**A**) Residue interaction energy (computed as the sum of the LJ and Coulombic interaction energies) comparison for active (red) and inactive (blue) AT1R conformational states. Data points show the interaction energy per residue (kJ/mol) averaged over the five MD simulation runs with standard deviation indicated by error bars. The interaction energy statistics are plotted in the order of AT1R residue number. Structural regions (transmembrane helices, intracellular loops, extracellular loops, and helix 8) are annotated and shaded to guide interpretation. (**B**) Violin plots summarizing interaction energy distributions across structural regions (N-terminal, TM1-7, ICL1-3, ECL1-3, and Helix 8) of active (red) and inactive (blue) receptor conformations. The median and quartile values are indicated within each distribution, revealing state-dependent differences in energy.

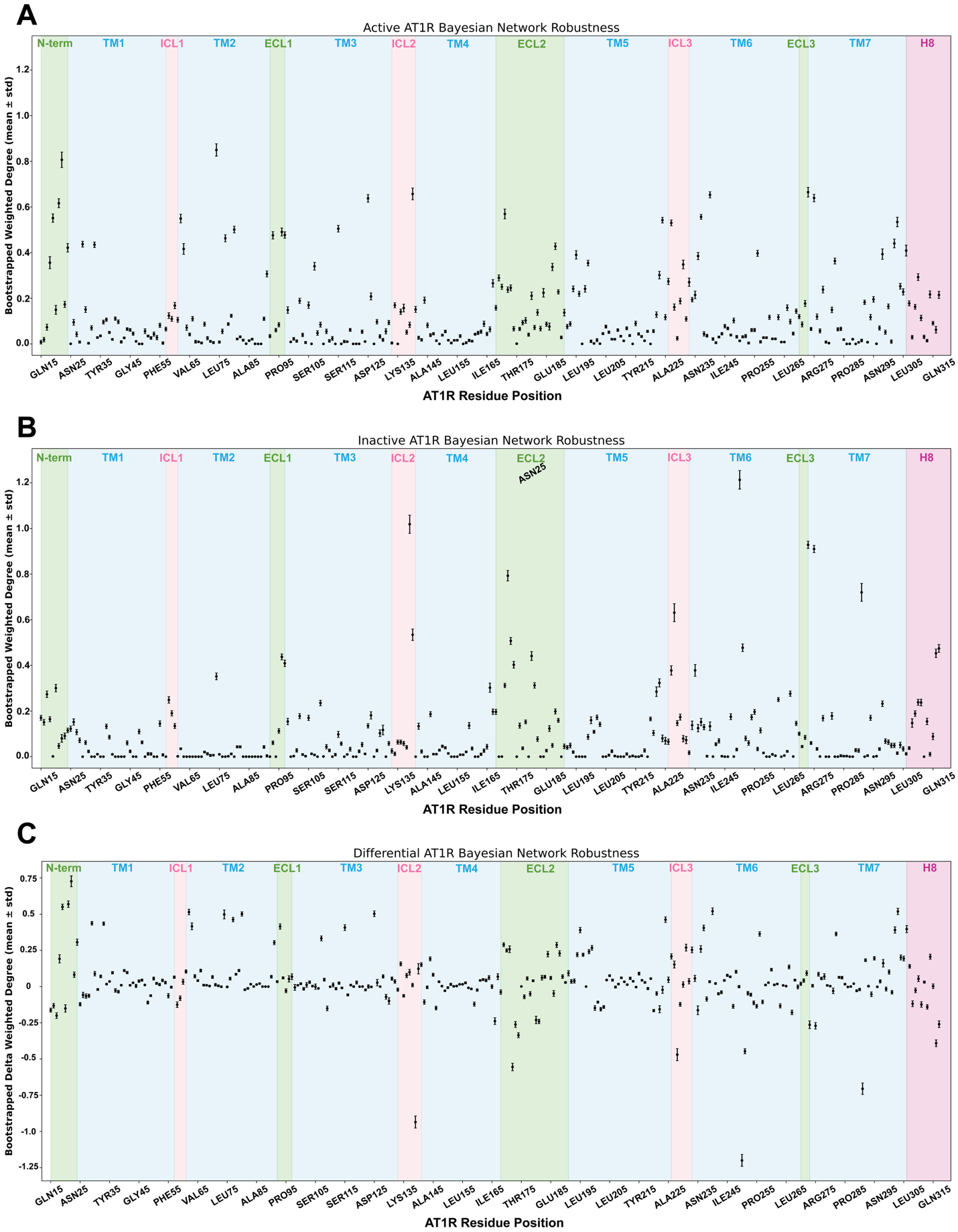

### Supplementary Fig. 2. Robustness of Bayesian Network edge weights and residue influence across 1,000 bootstrap resampling. (A) Weighted degree (Wd) robustness for residues in the active state. For each 1,000 bootstrap replicates, MI-based edge weights were recalculated from a resampled MD energy matrix and applied to the fixed BN topology. Points represent mean Wd per residue; vertical bars denote bootstrap standard deviations. Structural regions (transmembrane helices, intracellular loops, extracellular loops, and helix 8) are annotated and shaded to guide interpretation. (B) Corresponding Wd robustness analysis for the inactive, antagonist-bound AT1R ensemble. (C) Differential weighted degree (ΔWd) robustness between active and inactive states, computed as the difference between bootstrap-mean Wd values. Error bars denote propagated standard deviations from the two state-specific Wd distributions. Residues with consistently positive or negative ΔWd across replicates represent high-confidence state-dependent allosteric determinants. Together, these analyses demonstrate that MI-derived edge weights and node influence estimates are highly stable with respect to sampling variability in the MD dataset.

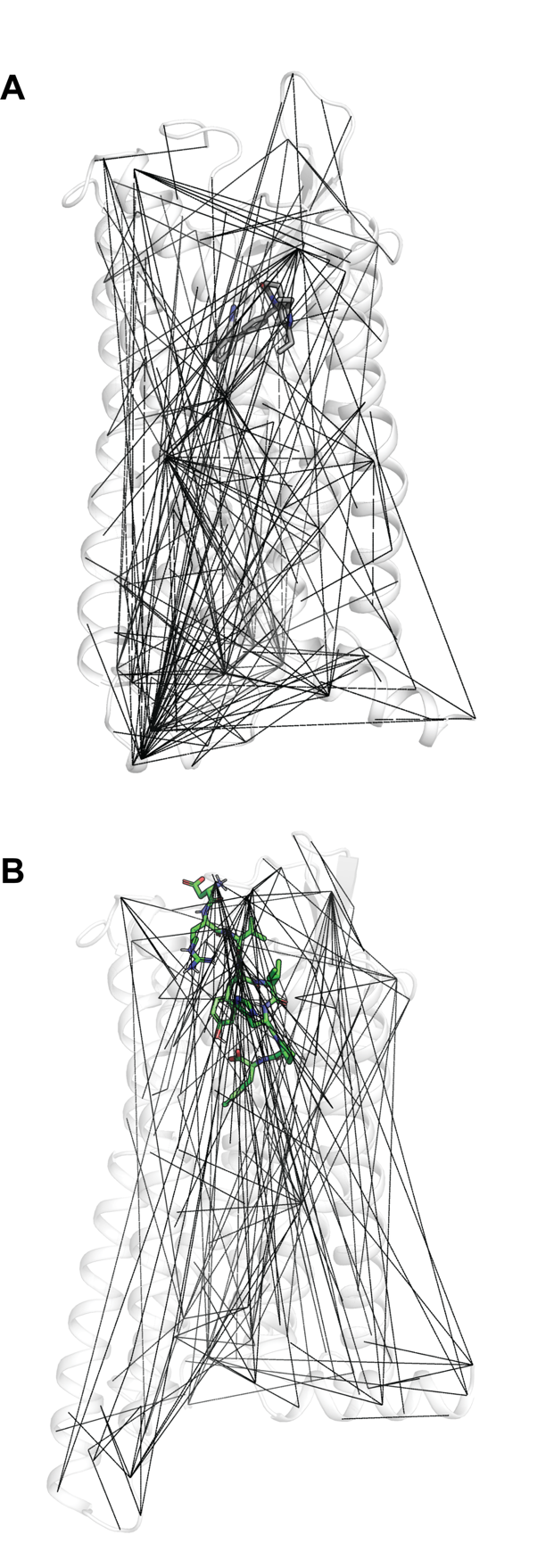

**Supplementary Fig. 3. Bayesian network (BN)–identified residue dependencies in AT1R.** Residue pairs identified as BN edges with separation >10 Å were mapped onto the (**A**) inactive AT1R (PDB ID: 4YAY) and (**B**) active AT1R (PDB ID: 7F6G) structures. Black lines represent BN-predicted residue dependencies that were supported by MI (mutual information) analysis (see Methods), indicating long-range dynamical couplings consistent with allosteric communication. These maps highlight the distinct distribution of long-range codependencies in the active versus inactive receptor conformations.

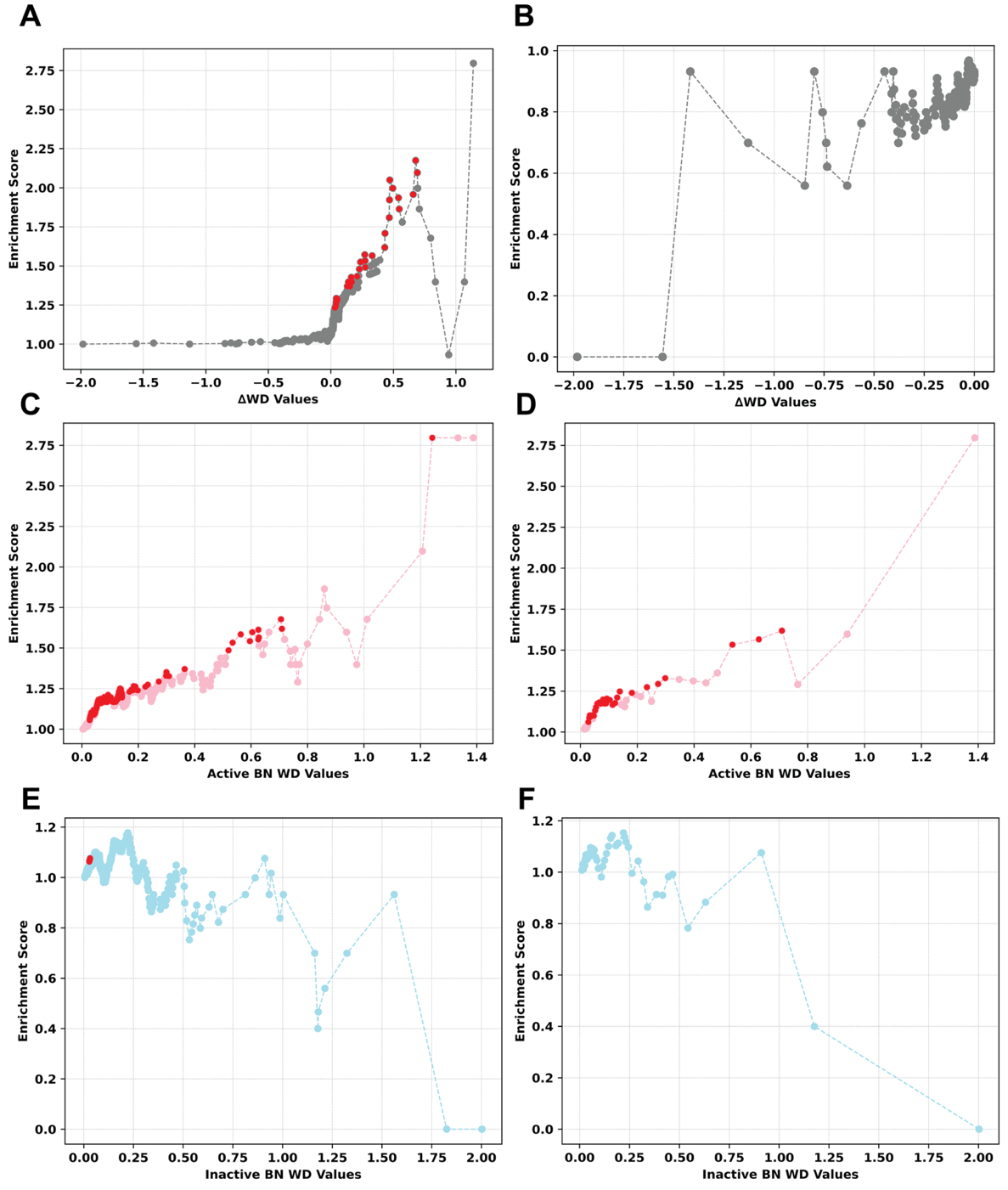

**Supplementary Fig. 4.** **Enrichment analysis of active- and inactive-state Bayesian Network (BN) predictions across residue-ranked thresholds.** (**A**) Full enrichment curve of residues ranked by ΔWD values. Each dot represents the enrichment score at a given *n* (top *n* residues ranked by ΔWD), comparing the proportion of true positives within the subset to the expected background rate. Red dots correspond to subset sizes that reach statistical significance (Fisher’s exact test *p* < 0.05), as shown in Fig. 2B. (**B**) Enrichment analysis of residues with negative ΔWD values (i.e., more connected in the inactive state than the active state). A significant number of bins fall below random expectation (enrichment score < 1), highlighting the model's predictive capacity on both ends of the dynamic range. (**C**) Enrichment analysis using WD values from the active-state BN only. Each dot represents an enrichment score at increasing *n* values. **(D)** Subsampled version of (C) showing every sixth dot to simplify visualization of trends. (**E**) Enrichment curve using WD values from the inactive-state BN**.** Enrichment scores remain near or below 1, indicating that high connectivity in the inactive-state BN is not predictive of functional importance. (**F**) Subsampled version of (E) showing every sixth dot for clarity.

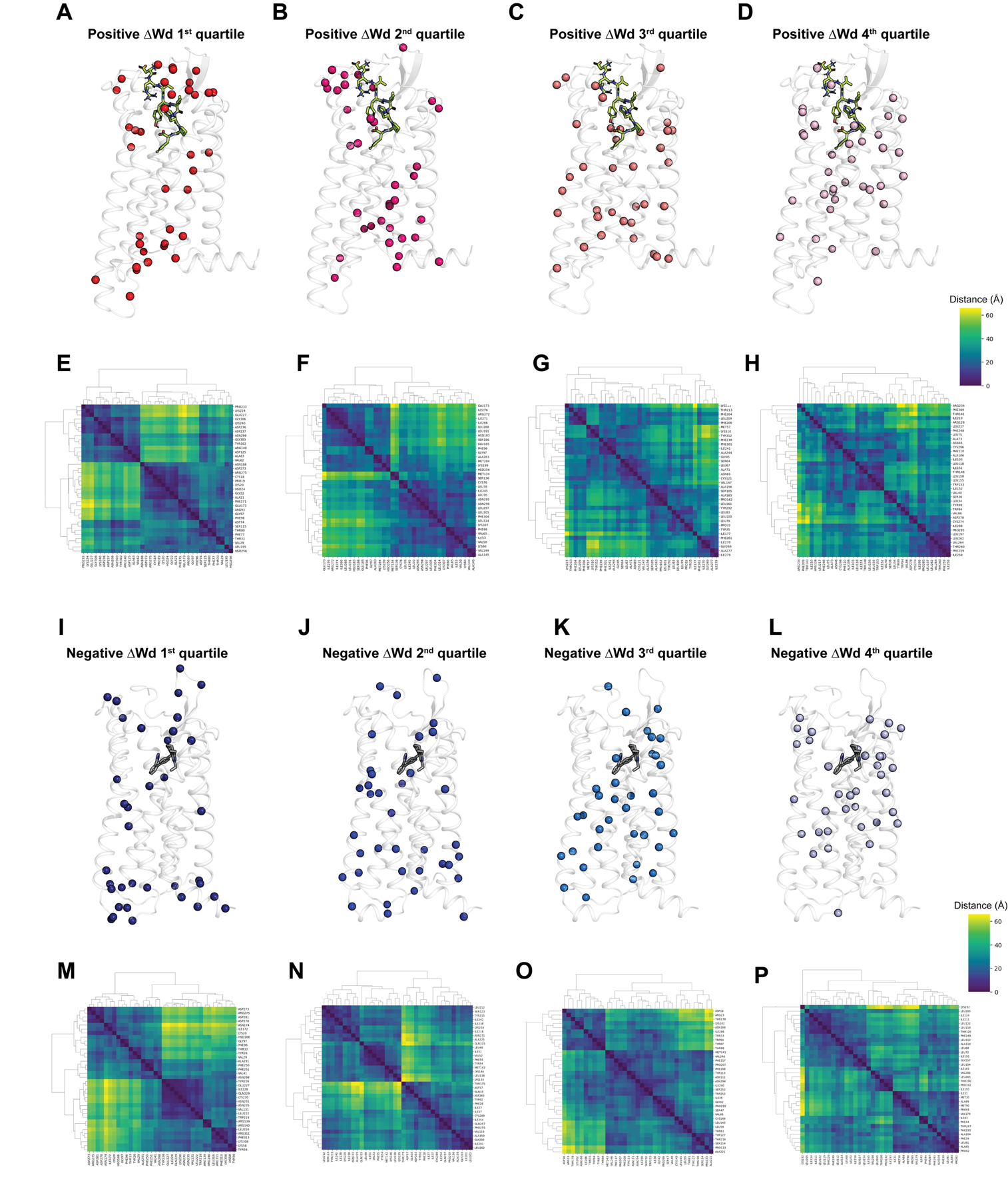

**Supplementary Fig. 5**. Quartile-based structural and clustering analysis of residues ranked by delta Weighted Degree (ΔWd). (**A**-**D**) Positive ΔWd quartiles (A, quartile 1 [highest positive]; B, quartile 2; C, quartile 3; D, quartile 4 [lowest positive]). Residues are mapped onto the active AT1R structure (PDB ID: 7F6G), with the (**E**-**H**) corresponding hierarchical clustering heatmaps below illustrating residue interaction patterns within each quartile. (**I**-**L**) Negative ΔWd quartiles (I, quartile 1 [most negative]; J, quartile 2; K, quartile 3; L, quartile 4 [least negative]). Residues are mapped onto the inactive AT1R structure (PDB ID: 4YAY), alongside (**M**-**P**) their respective hierarchical clustering heatmaps. Residue colors in structural views correspond to quartile strength, with darker colors indicating stronger differential degrees (more significant contributions to active or inactive network differences). Heatmaps show hierarchical clustering based on interaction similarity within each quartile, highlighting distinct structural and functional patterns among residue groups.

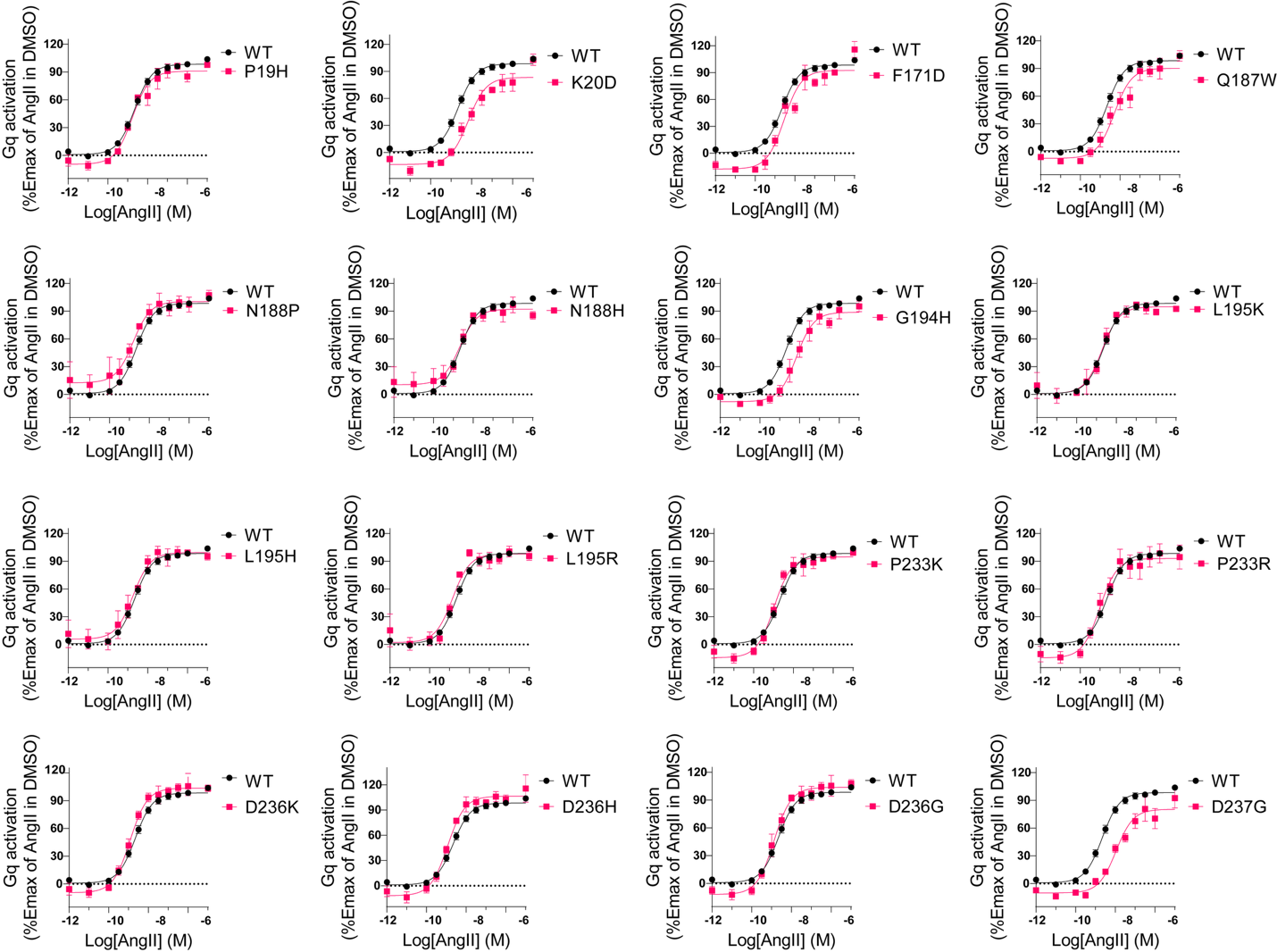

**Supplementary Fig. 6.** **Dose response curves for AngII-mediated Gq signaling on WT and non-alanine mutants of AT1R.** HEK293 cells in 96-well plates were transfected with AT1R WT or mutant DNA along with Gq sensor (Gα_q_, P63-RlucII and rGFP-caax). Cells were then stimulated with various concentrations of AngII for 3 min before BRET measurement. Data are from at least three independent experiments and expressed as mean ± SEM.

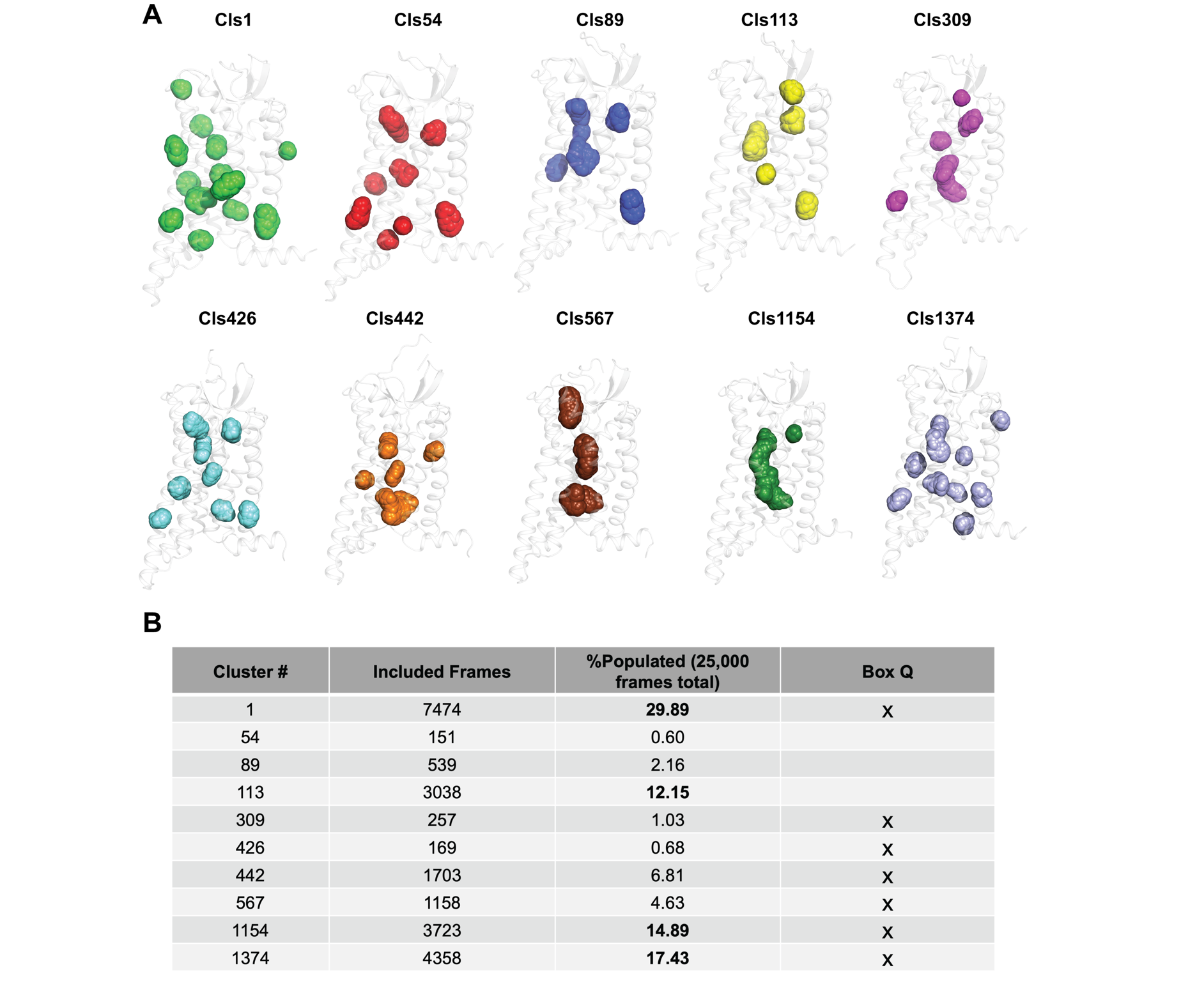

**Supplementary Fig. 7. Conformational clusters of AT1R identified by RMSD-based clustering of active MD trajectories.** (**A**) Representative structures of the top ten conformational clusters obtained from RMSD-based clustering of the MD simulation trajectories of the active state of AT1R (see Methods). Each cluster conformation is shown with putative small molecule binding pockets in distinct colors corresponding to the specific cluster ID. (**B**) Summary of clustering statistics, including the number of frames assigned to each cluster, percent population, and pocket population. The most populated clusters (Cls1, Cls113, Cls1154, Cls1374) together represent >70% of the sampled conformational ensemble, indicating dominant conformational states of AT1R captured during the clustering.

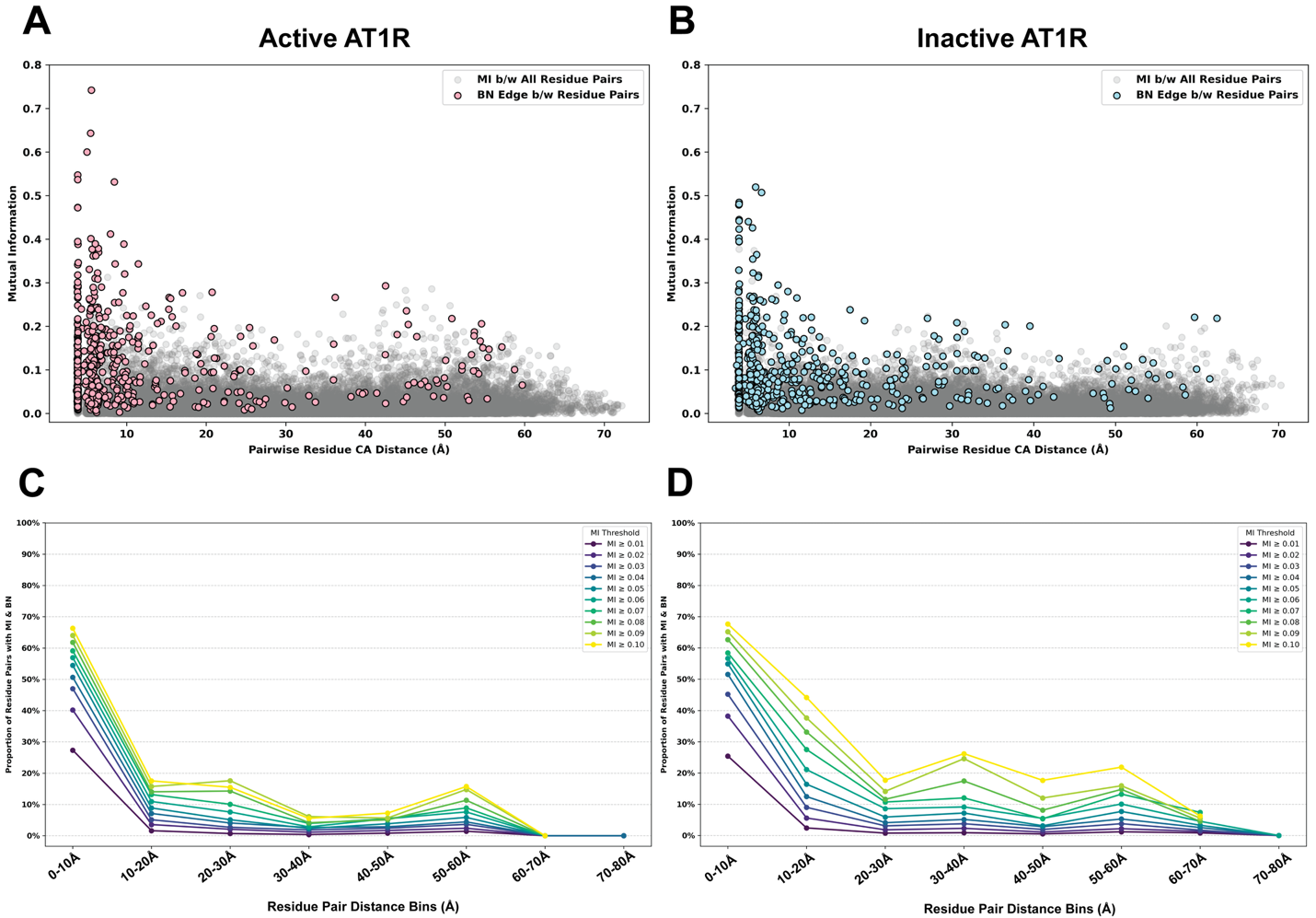

**Supplementary Fig. 8. Analysis of residue pair mutual information (MI) and Bayesian network (BN) edges in active and inactive AT1R.** (**A**-**B**) Scatter plots displaying all pairwise residue mutual information values (gray dots) in (**A**) active and (**B**) inactive AT1R systems based on their physical distances. Residue pairs connected by BN edges in each state-specific Bayesian network are highlighted (red circles for active, blue circles for inactive). **(C-D)** Proportion of BN-connected residue pairs at varying MI thresholds plotted against pairwise residue distances. Both active (**C**) and inactive (**D**) systems exhibit prominent enrichment of BN edges at short residue distances (<10 Å) and a secondary enrichment at longer, allosteric distances (>30 Å).

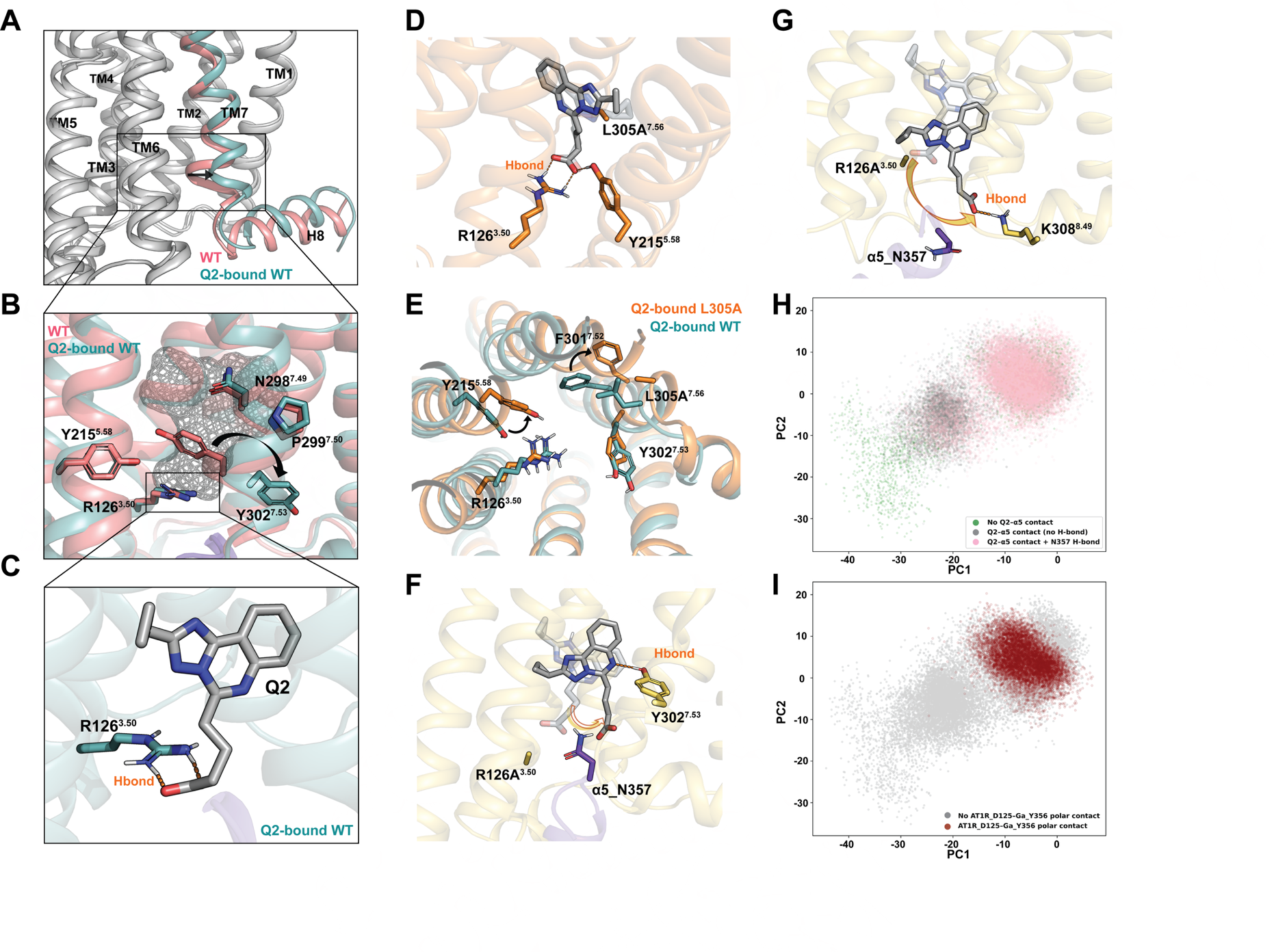
 **Supplementary Fig. 9. Molecular mechanisms underlying mutation-enhanced negative allosteric modulation by Q2.** (**A**) Overall view of the intracellular region of AT1R comparing AngII-bound WT (pink) and Q2-bound WT (cyan) with only TM7 colored. (**B**) Superposition of Q2-bound WT AT1R (cyan) and WT AT1R without Q2 (pink) highlighting the intracellular allosteric pocket adjacent to the G protein interface. NPxxY and selected microswitch residues lining the pocket are shown as sticks. The mesh represents the Q2 occupancy volume. (**C**) Close-up view of the WT Q2-binding pose showing a stabilizing hydrogen bond between Q2 and R126³·⁵⁰ that anchors the ligand above R126 in the upper region of the intracellular pocket. (**D**) Representative structure from MD simulations of the L305A⁷·⁵⁶ mutant showing bifurcated hydrogen bonds of Q2 with both R126³·⁵⁰ and Y215⁵·⁵⁸. (**E**) Structural overlay of Q2-bound WT (cyan) and Q2 bound L305A⁷·⁵⁶ (orange) AT1R. (**F**) Representative snapshot from MD simulations of the R126A³·⁵⁰ mutant showing loss of the R126³·⁵⁰–Q2 hydrogen bond and increased ligand mobility. Q2 forms alternative stabilizing interactions with Y302⁷·⁵³. (**G**) In the R126A³·⁵⁰ mutant, Q2 makes hydrogen bond K308⁸·⁴⁹, positioning the ligand closer Gα_q_ C-terminal helix (Gα_q__N357 shown in purple). (**H**) Projection of α5 helix conformations from Q2-bound R126A³·⁵⁰ simulations onto the principal component (PC) space, colored by the presence or absence of Q2–α5_N357 contact without hydrogen bonding. (**I**) Same PC projection as in (H), colored by the presence of polar contacts between AT1R D125³·⁴⁹ and Gα_q__Y356.

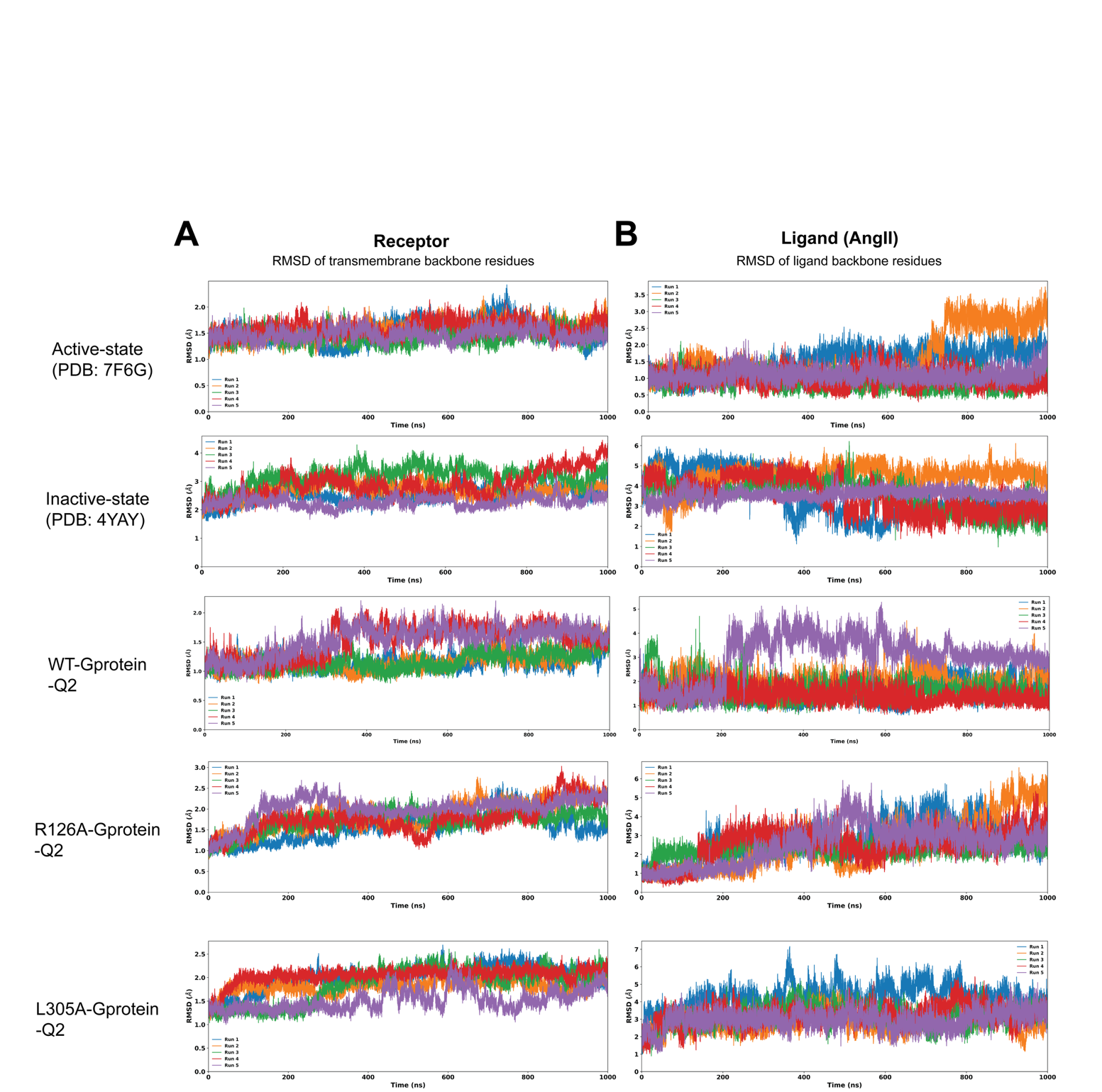
 **Supplementary Fig. 10. Analysis for convergence of MD simulation trajectories.** (**A**) Root-mean-square deviation (RMSD) of transmembrane backbone residues for 5 independent runs in active- (with G protein), inactive-states and 3 mutants (with G protein and Q2). (**B**) RMSD of ligand backbone residues for 5 independent runs in active- (with G protein), inactive-states and 3 mutants (with G protein and Q2).

**Supplementary Table 1. Pharmacological parameters of AngII-mediated Gq signaling on AT1R WT and mutants.** WT and mutant receptors were expressed in HEK 293 cells with Gq sensor (Gα_q_, P63-RlucII and rGFP-caax). BRET assay was used to measure the AngII-mediated Gα_q_ activation. Data are normalized to the Emax of the WT receptor. The pEC_50_, Emax, and Relative activity (RA) were obtained from the data in Fig.4B. Emax is reported as the difference between the basal and top values. RA (Δ(log(Emax/EC_50_))) was calculated by subtracting the WT log(Emax/EC_50_) from the mutant log(Emax/EC_50_). *Indicating statistically significant difference compared to the WT receptor (p < 0.05), obtained with unpaired Student’s t-test. Data are the mean ± SEM of at least three independent experiments.

| **AT1R Mutation** | pEC_50_  (CL95%) | Emax ± SEM | RA ΔLog(Emax/EC_50_) | Expression (%WT) |
| --- | --- | --- | --- | --- |
| **WT** | 8.65 (8.57 - 8.74) | 100 |  | 100 |
| **P19H** | 8.77 (8.59 - 8.95) | 100.6 ± 4.2 | -0.12 ± 0.07 | 102 |
| **K20D** | 8.17 (7.93 - 8.4) | 96.7 ± 4.5 | 0.06 ± 0.18 | 107 |
| **A21W** | 6.34^*^ (6.19 - 6.49) | 76.6 ± 3.6^*^ | -2.39 ± 0.14^*^ | 87 |
| **H24P** | 7.69^*^ (7.45 - 7.95) | 100.5 ± 4.7 | -0.89 ± 0.24^*^ | 144 |
| **S107P** | 6.38^*^ (5.56 - 7.21) | 15 ± 3.2^*^ | -2.36 ± 0.57^*^ | 60 |
| **V169W** | 7.49^*^ (7.31 - 7.68) | 94.7 ± 3.9 | -0.88 ± 0.21^*^ | 82 |
| **F171D** | 8.54 (8.27 - 8.8) | 110.2 ± 5.9 | -0.14 ± 0.17 | 143 |
| **Q187W** | 8.29 (8 - 8.57) | 97.5 ± 5.2 | -0.07 ± 0.19 | 135 |
| **N188I** | 8.81 (8.68 - 8.95) | 92.4 ± 2.9^*^ | -0.15 ± 0.12 | 108 |
| **N188H** | 8.67 (8.39 - 8.97) | 81.7 ± 5.8^*^ | -0.36 ± 0.3 | 112 |
| **N188P** | 8.81 (8.47 - 9.16) | 87.8 ± 7.1 | -0.17 ± 0.11 | 122 |
| **G194H** | 8.15 (7.93 - 8.38) | 96.7 ± 4.8 | -0.19 ± 0.2 | 125 |
| **L195E** | 7.81^*^ (7.65 - 7.98) | 91.9 ± 3.5^*^ | -1.15 ± 0.1^*^ | 107 |
| **L195K** | 8.73^*^ (8.52 - 8.93) | 93.9 ± 4.7^*^ | -0.21 ± 0.03^*^ | 90 |
| **L195H** | 8.74 (8.48 - 9.01) | 93.7 ± 5.8 | -0.18 ± 0.02^*^ | 95 |
| **L195R** | 8.84 (8.63 - 9.06) | 95.8 ± 5.6 | -0.09 ± 0.11 | 104 |
| **T198H** | 8.97 (8.79 - 9.15) | 113.9 ± 4.8^*^ | 0.29 ± 0.16 | 112 |
| **P233K** | 8.98 (8.82 - 9.13) | 109 ± 4.3 | 0.11 ± 0.11 | 95 |
| **P233L** | 8.72 (8.59 - 8.86) | 91 ± 3.1^*^ | -0.23 ± 0.07^*^ | 115 |
| **P233R** | 9.02 (8.75 - 9.28) | 107.5 ± 6.7 | 0.15 ± 0.19 | 124 |
| **D236G** | 8.88 (8.71 - 9.05) | 115.7 ± 4.5 | 0.07 ± 0.12 | 62 |
| **D236R** | 9.02 (8.88 - 9.16) | 116.2 ± 3.9^*^ | 0.18 ± 0.1 | 102 |
| **D236H** | 8.94 (8.77 - 9.1) | 118.4 ± 4.7 | 0.1 ± 0.11 | 90 |
| **D236K** | 8.92 (8.78 - 9.07) | 113.1 ± 3.8 | 0.07 ± 0.05 | 86 |
| **D237G** | 7.96 (7.71 - 8.22) | 90.4 ± 4.7 | -0.18 ± 0.15 | 94 |
| **K240P** | 8.95 (8.74 - 9.15) | 106.8 ± 5.1^*^ | 0.24 ± 0.15 | 98 |

**Supplementary Table 2. Structures of Q compounds and their effect on AngII-mediated Gq activation.** HEK293 cells expressing AT1R and the Gq BRET biosensor were pretreated with 200 μM of Q compounds before AngII stimulation.

| **Name** | **Structure** | | **Luciferase activity** | | **Emax** | | **ΔLogEC_50_** |
| --- | --- | --- | --- | --- | --- | --- | --- |
|  | **SMILES** | | (%DMSO) | | mean ± SEM | | mean ± SEM |
| **Q1** | 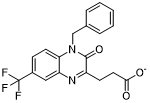 | 81 | | 110.1 ± 1.9 | | 0.057 ± 0.044 | |
|  | [O-]C(=O)CCc(n1)c(=O)n(c(c12)ccc(C(F)(F)F)c2)Cc3ccccc3 |  |  |  |  |  |  |
| **Q2** | 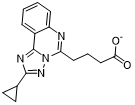 | 101 | | 100.1 ± 1.7 | | 0.325 ± 0.052 | |
|  | c1cccc(c1c23)nc(CCCC([O-])=O)n3nc(n2)C4CC4 |  |  |  |  |  |  |
| **Q3** | 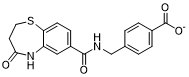 | 106 | | 101.8 ± 2.7 | | 0.089 ± 0.074 | |
|  | [O-]C(=O)c1ccc(cc1)CNC(=O)c(c2)ccc(c23)SCCC(=O)N3 |  |  |  |  |  |  |
| **Q4** | 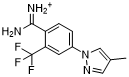 | 71 | | 106.1 ± 2.7 | | 0.035 ± 0.076 | |
|  | NC(=[NH2+])c1c(C(F)(F)F)cc(cc1)-n(c2)ncc2C |  |  |  |  |  |  |
| **Q5** | 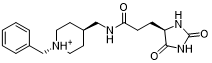 | 99 | | 106.8 ± 2.0 | | 0.038 ± 0.052 | |
|  | N1C(=O)N[C@H](C1=O)CCC(=O)NC[C@H]2CC[N@@H+](CC2)Cc3ccccc3 |  |  |  |  |  |  |
| **Q6** | 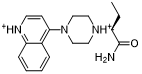 | 91 | | 106.9 ± 2.0 | | 0.054 ± 0.023 | |
|  | CC[C@@H](C(=O)N)[NH+](CC1)CCN1c2cc[nH+]c(c23)cccc3 |  |  |  |  |  |  |
| **Q7** | 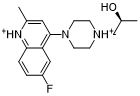 | 82 | | 108.1 ± 2.4 | | -0.027 ± 0.051 | |
|  | C[C@H](O)C[NH+](CC1)CCN1c2cc(C)[nH+]c(c23)ccc(F)c3 |  |  |  |  |  |  |
| **Q8** | 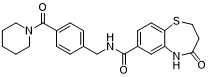 | 50 | | 125.5 ± 3.2 | | 0.131 ± 0.018 | |
|  | N1C(=O)CCSc(c12)ccc(c2)C(=O)NCc(cc3)ccc3C(=O)N4CCCCC4 |  |  |  |  |  |  |
| **Q9** | 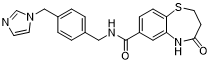 | 63 | | 128.0 ± 2.4 | | 0.104 ± 0.089 | |
|  | N1C(=O)CCSc(c12)ccc(c2)C(=O)NCc3ccc(cc3)Cn4ccnc4 |  |  |  |  |  |  |

| **Table S3. AngII potencies of G_q/11_ pathways on wild-type (WT) and mutant AT1Rs in the absence (DMSO) or the presence of Q2.** WT and mutant receptors were expressed in HEK 293 cells along with G_q/11_ sensor (P63-RlucII and rGFP-caax) to measure G_q/11_ activities. Cells were preincubated with 200 μM of Q2 for 30 min before AngII stimulations. Data are normalized to Emax of WT in the absence of Q2 (DMSO). The basal, Emax, and LogEC_50_ in nM were obtained from the data presented in Fig.7. Each value represents mean ± SEM of more than three independent experiments. LogEC_50_ changes (ΔLogEC_50_) were calculated by subtracting LogEC_50_ value in the absence of Q2 (DMSO) from LogEC_50_ in the presence of 200 μM Q2 in each experiment and averaged. ^*, #^ p < 0.05, ^**, ##^ p < 0.01, ^***, ###^ p < 0.001, ^****, ####^ p < 0.0001, unpaired Student’s t-test compared to the value obtained in the absence of Q2 (DMSO) (^*^) or compared to the value of WT (^#^). |  | |  | |  |  | |  | |  |  |  |  |  |
| --- | --- | --- | --- | --- | --- | --- | --- | --- | --- | --- | --- | --- | --- | --- |
| BW |  | |  | |  | | Basal | |  | Emax |  | LogEC_50_ |  | ΔLogEC_50_ |
|  |  | |  | |  | |  |  |  |  |  |  |  | (Q2-DMSO) |
|  | WT | | | DMSO |  | | 0.2 ± 1.1 | |  | 99.9 ± 0.9 |  | -8.75 ± 0.03 |  | 0.50 ± 0.03 |
|  |  |  |  | Q2 |  | | -1.9 ± 1.6 | |  | 100.7 ± 1.7 |  | -8.23 ± 0.04^****^ |  |  |
| 2.46 | | L70A | | DMSO |  | | 2.8 ± 4.4 | |  | 74.3 ± 3.8 |  | -8.54 ± 0.07 |  | 0.53 ± 0.14 |
|  |  |  |  | Q2 |  | | -1.4 ± 3.8 | |  | 58.0 ± 4.2* |  | -8.01 ± 0.13^*^ |  |  |
| 2.50 | | D74A | | DMSO |  | | -1.4 ± 1.2 | |  | 17.6 ± 1.9 |  | -7.50 ± 0.27 |  | 0.69 ± 0.37 |
|  |  |  |  | Q2 |  | | -3.1 ± 1.0 | |  | 23.1 ± 2.2 |  | -6.81 ± 0.11^*^ |  |  |
| 3.43 | | L119A | | DMSO |  | | 2.0 ± 2.4 | |  | 67.5 ± 1.7 |  | -8.54 ± 0.02 |  | 0.74 ± 0.11^#^ |
|  |  |  |  | Q2 |  | | -1.7 ± 2.2 | |  | 64.3 ± 2.8 |  | -7.80 ± 0.13^**^ |  |  |
| 3.50 | | R126A | | DMSO |  | | 7.8 ± 2.8 | |  | 62.6 ± 3.0 |  | -8.09 ± 0.10 |  | 0.87 ± 0.07^##^ |
|  |  |  |  | Q2 |  | | 6.3 ± 2.3 | |  | 64.6 ± 4.1 |  | -7.21 ± 0.10^***^ |  |  |
| 6.41 | | V246A | | DMSO |  | | -0.7 ± 2.2 | |  | 64.2 ± 2.2 |  | -8.29 ± 0.04 |  | 0.49 ± 0.11 |
|  |  |  |  | Q2 |  | | -1.9 ± 2.8 | |  | 69.5 ± 3.5 |  | -7.80 ± 0.14^*^ |  |  |
| 6.43 | | F248A | | DMSO |  | | 6.6 ± 3.1 | |  | 92.1 ± 2.6 |  | -8.61 ± 0.05 |  | 0.03 ± 0.12^###^ |
|  |  |  |  | Q2 |  | | -0.1 ± 4.2 | |  | 97.3 ± 3.6 |  | -8.58 ± 0.11 |  |  |
| 7.45 | | N294A | | DMSO |  | | 0.6 ± 1.6 | |  | 32.2 ± 1.8 |  | -7.98 ± 0.12 |  | 0.36 ± 0.11 |
|  |  |  |  | Q2 |  | | -3.4 ± 1.4 | |  | 23.9 ± 1.8* |  | -7.62 ± 0.13 |  |  |
| 7.46 | | N295A | | DMSO |  | | 6.6 ± 3.4 | |  | 101.4 ± 3.6 |  | -8.14 ± 0.02 |  | 0.13 ± 0.12^##^ |
|  |  |  |  | Q2 |  | | 5.4 ± 3.9 | |  | 100.2 ± 4.4 |  | -8.01 ± 0.13 |  |  |
| 7.49 | | N298A | | DMSO |  | | -1.0 ± 1.9 | |  | 59.3 ± 2.0 |  | -8.15 ± 0.06 |  | 0.34 ± 0.17 |
|  |  |  |  | Q2 |  | | 0.1 ± 1.8 | |  | 47.3 ± 2.2* |  | -7.81 ± 0.14 |  |  |
| 7.50 | | P299A | | DMSO |  | | 3.4 ± 3.5 | |  | 54.4 ± 3.2 |  | -8.41 ± 0.06 |  | 0.73 ± 0.07^##^ |
|  |  |  |  | Q2 |  | | 4.7 ± 3.5 | |  | 63.4 ± 4.6 |  | -7.68 ± 0.10^**^ |  |  |
| 7.52 | | F301A | | DMSO |  | | 4.4 ± 1.9 | |  | 48.1 ± 2.6 |  | -7.68 ± 0.05 |  | -0.01 ± 0.07^####^ |
|  |  |  |  | Q2 |  | | 2.3 ± 3.0 | |  | 45.5 ± 4.0 |  | -7.69 ± 0.10 |  |  |
| 7.53 | | Y302A | | DMSO |  | | -1.7 ± 2.7 | |  | 61.0 ± 2.7 |  | -8.19 ± 0.08 |  | 0.57 ± 0.06 |
|  |  |  |  | Q2 |  | | -3.1 ± 2.8 | |  | 66.6 ± 3.9 |  | -7.62 ± 0.03^**^ |  |  |
| 7.56 | | L305A | | DMSO |  | | 4.0 ± 2.6 | |  | 101.7 ± 2.1 |  | -8.73 ± 0.04 |  | 0.89 ± 0.04^####^ |
|  |  |  |  | Q2 |  | | 3.1 ± 2.4 | |  | 102.3 ± 3.0 |  | -7.84 ± 0.08^****^ |  |  |

**Supplementary Data 1. Bayesian Network Metrics Weighted Degree (For Fig.2)**

**Supplementary Data 2. Deep Mutational Scanning data of all Ala-null top quartile residues**

**Supplementary Data 3. Primer sequences for generation of AT1R mutants**

**Supplementary Network files 1-4. Bayesian Network Files (.graphml files to be opened in Cytoscape/Gephi) (For Fig.3)**
